# Molecular cloning of a novel, nervous system-specific RGS6 isoform lacking canonical G protein regulatory effects and with dominant negative actions

**DOI:** 10.64898/2026.05.08.723811

**Authors:** KE Ahlers-Dannen, J Yang, J Bernholtz, Alexander Glebov-Mccloud, Stefan Strack, JG Koland, RA Fisher, A Stewart

## Abstract

Regulator of G protein Signaling 6 (RGS6), heavily implicated in neurological and neuropsychiatric disorders, is enriched in mouse and human brain. Our initial cloning effort identified 36 RGS6 mRNAs in human brain. However, we recently identified an additional RGS6 protein isoform that is larger (∼69kDa) than the ubiquitously expressed ∼56kDa RGS6L(+GGL) isoforms. Notably, this isoform, named “RGS6B” for “brain-specific”, is selectively expressed in the nervous system of mice and humans. Here, we report the cloning of a new RGS6-encoding mRNA, which resembles the RGS6Lα1(+GGL) transcript identified in our initial cloning effort but includes a highly conserved novel exon (Alternative 3, A3) that alters the reading frame of terminal exon α resulting in an extension of the protein C-terminus. When expressed in cells, RGS6LA3α1(+GGL) co-migrates with RGS6B, and, importantly, interfering RNA targeting exon A3 results in selective depletion of RGS6B in isolated primary cortical astrocytes. RGS6B is capable of stabilizing RGS6 binding partners R7BP and Gβ_5_ and, in fact, exhibits an increased protein half-life relative to RGS6L. Both RGS6L and RGS6B are downregulated in human gliomas and share the ability to kill U87MG glioblastoma cells when overexpressed indicating conservation of non-canonical cytotoxic activity between RGS6L and RGS6B species. However, RGS6B lacks the ability to counteract Gα_i/o_-dependent suppression of cAMP signaling, indicating a lack of functional GTPase activating protein (GAP) activity. Instead, RGS6B functions in a dominant negative manner to block Gα_i/o_ regulation by RGS6L. RGSB is the first identified RGS protein member that functions to promote, rather than inhibit, G protein signaling. The discovery of the molecular identity of RGS6B will now allow for delineation of unique functions for RGS6 protein isoforms in both physiological and pathophysiological brain states.

## Introduction

In 2019, seminal work conducted by the Cross-Disorder Group of the Psychiatric Genomics Consortium identified a single nucleotide polymorphism (SNP) in *RGS6* (rs2332700) as one of only 23 loci linked to ≥4 complex neuropsychiatric disorders at genome wide significance. This *RGS6* SNP was strongly associated with major depressive disorder (MDD), bipolar disorder (BPD), autism spectrum disorder (ASD), and schizophrenia.^1^ Notably, the Genome Wide Association Study (GWAS) Catalog^2^ robustly supports and extends this finding, associating *RGS6* variants with not only psychiatric disorders (e.g., schizophrenia, substance use disorders, MDD), but also neurodegenerative disorders (e.g., Alzheimer’s disease), cognitive function, brain malignancies, and more (**Fig. 1A**). Interestingly, regulator of G protein signaling 6 (RGS6) is the most frequent RGS family member represented in the GWAS Catalog in these categories, even among other highly homologous R7 subfamily members (RGS’s 6, 7, 9, and 11) (**Fig. 1A**).

**Figure 1.**
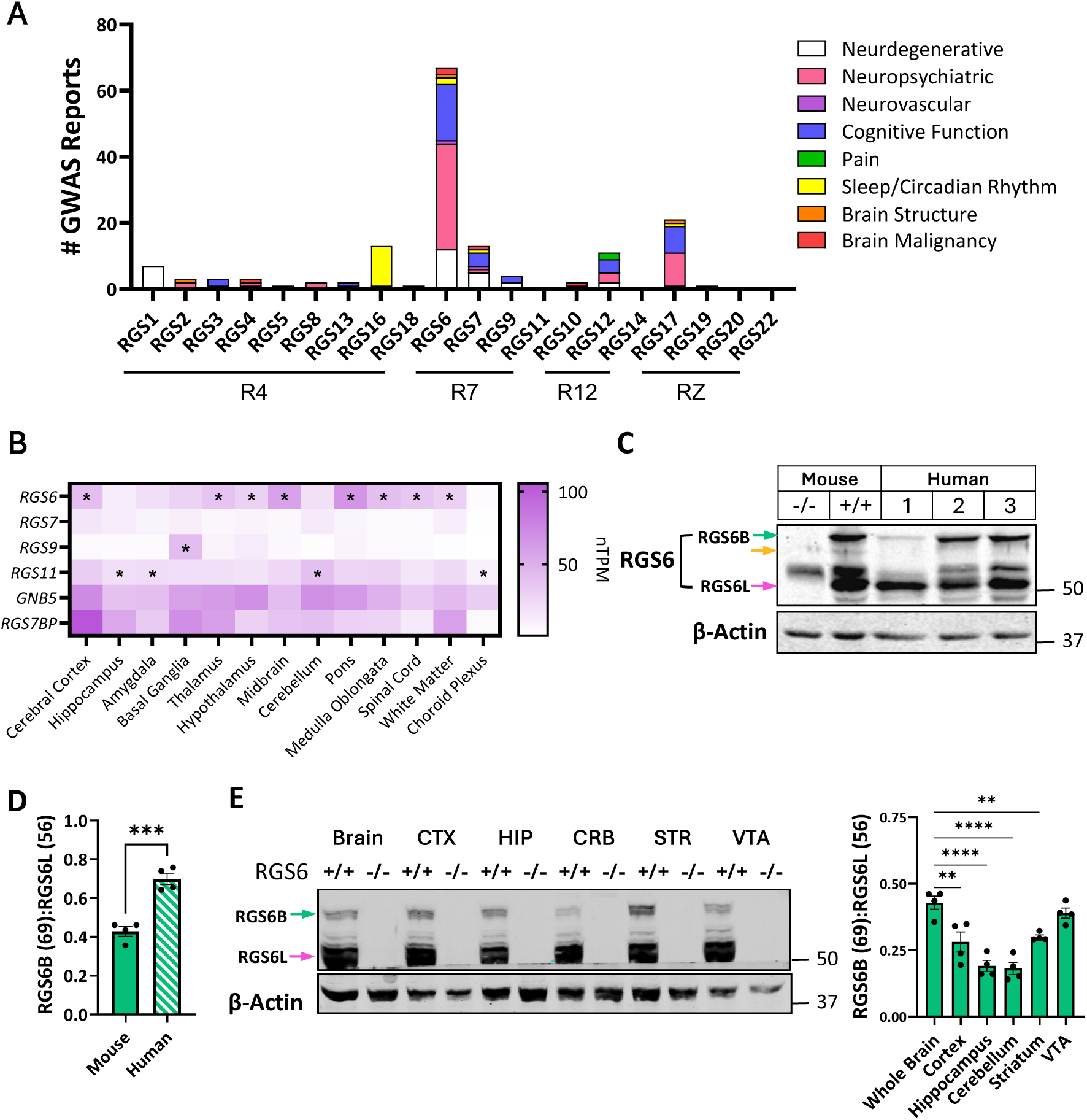
The RGS6B protein is conserved between humans and mice and is expressed in brain regions implicated in neurological disorders . (**A**) Tabulation of GWAS reports indexed on GWAS Catalog linking members of the RGS superfamily to brain function and health. In total, we identified 67 studies linking RGS6 to nervous system function or dysfunction. For studies seeking to elucidate the genetic underpinnings of neurodegenerative disorders, neuropsychiatric conditions, cognitive function, and brain cancers, RGS6 is the most implicated RGS protein. (**B**) mRNA expression of R7 family RGS proteins as well as their Gβ5 and R7BP binding partners across subregions of the human brain. Data are expressed as transcripts per million (TPM) normalized across multiple source datasets available on the Human Protein Atlas. A * denotes R7 family member with highest regional mRNA expression. (**C**) Immunoblotting of RGS6L protein expression in whole brain from human and WT or RGS6-/- mice. (**D**) The ratio of RGS6B (69 kDa) to RGS6L(+GGL) (56 kDa) was determined via densitometric analysis of immunoblots (n=4). (**E**) RGS6L(+GGL) protein expression across brain subregions isolated from WT or RGS6-/- mice. The ratio of RGS6B (69 kDa) to RGS6L(+GGL) (56 kDa) was determined via densitometric analysis of immunoblots (n=4). Data were analyzed via Student’s t-test or one-way ANOVA with Dunnett’s post-hoc test. **P<0.01, ***P<0.001, ****P<0.0001. Data are expressed as mean ± S.E.M.

Canonically, RGS proteins modulate G protein-coupled receptor (GPCR) signaling by facilitating heterotrimeric G-protein inactivation, a function bestowed by the GTPase-activating activity of their RGS domain towards Gα subunits. R7 family RGS proteins share 2 additional domains. First, the N-terminal disheveled, EGL-10, pleckstrin homology (DEP)/DEP helical extension (DHEX) domain is best known for mediating interaction with an accessory protein, R7 family-binding protein (R7BP), which facilitates membrane localization of R7 family members and, given its expression is confined to the central nervous system^3^, augments R7 RGS-dependent GPCR blockade in neurons.^4,5^ Second, the G gamma subunit-like (GGL) domain, located between the N-terminal DEP/DHEX domains and the C-terminal RGS domain, facilitates binding between R7 family members and the atypical Gβ subunit Gβ_5_, an interaction required for the stability of both Gβ_5_ and R7 family RGS proteins.^6^

A review of the Human Protein Atlas indicates that RGS6 mRNA expression is highest in the brain, that it is detectable throughout the central nervous system (CNS), and that it is the most highly expressed R7 family member in most regions surveyed (**Fig. 1B**). RGS6 is highly pleiotropic having been previously implicated in motor coordination^7,8^, mood^9^, learning and memory^10^, and alcohol consumption^11,12^ via its ability to promote neurogenesis^10^, ensure neuroprotection^8^, and modulate the activity of Gα_i/o_-coupled GABA B receptors (GABA_B_Rs)^7^, serotonin (5-HT) 1A receptors (5-HT_1A_Rs)^9,13^ and dopamine (DA) D2 receptors (D2Rs).^11,14^ In non-neuronal cell types, RGS6 also possess potent pro-apoptotic^15–19^ and growth suppressive^20^ capabilities and has been identified as a tumor suppressor in breast and bladder.^21,22^ Efforts to elucidate the mechanistic role of RGS6 in disease pathogenesis have transitioned from study of global RGS6 knockout (RGS6^-/-^) models to approaches that allow for improved spatial and temporal control of gene manipulation (e.g., RGS6 floxed mice, RGS6^flx/flx^).^10,12^ However, complex alternate splicing of the RGS6 gene adds an additional layer of complexity to the elucidation of RGS6 biology. 36 distinct RGS6 splice variants have been identified which are predicted to produce viable RGS6 protein isoforms containing either long (RGS6L) or short (RGS6S) N-terminal domains, an incomplete or intact GGL domain (-/+ GGL), and nine alternative C-terminal sequences.^23^ Identification of an RGS6 variant that disrupts gene splicing (1369-1 G>C) associated with human cataracts, mental retardation, and microcephaly^24^ alludes to functional importance of RGS6 alternative splicing in the CNS, but the predominant RGS6 isoforms expressed across tissue/organ systems remain unclear.

Recently, progress has been made on this front as we developed three distinct antibodies against the whole RGS6L protein (RGS6-fl), the N-terminus of the RGS6L isoforms (RGS6-L), and an alternative amino acid sequence found in 56% of known RGS6 isoforms (RGS6-18). Using these antibodies, we defined, in mouse, whole-body tissue-specific differences in RGS6 isoform expression.^25^ Importantly, in that study, we identified novel highly conserved RGS6 protein bands corresponding to phosphorylated and dephosphorylated forms of a nervous system specific RGS6 isoform (RGS6 “Brain”; RGS6B),^25^ which we hypothesize plays an important role in brain health. Of particular interest here, reexamination of western blot data presented in our previous work revealed that, in addition to the ubiquitously expressed RGS6L(+GGL) isoforms, RGS6B is also highly expressed in prefrontal cortex, hippocampus, cerebellum, striatum, nucleus accumbens, and ventral tegmental area (VTA). Here, we expand the RGS6 splicing scheme to include 9 additional exons yielding a possible 420 unique RGS6 splice forms. Most notably, we report cloning and initial functional characterization of a new RGS6 transcript, arising via inclusion of a novel exon (Alternative 3, A3) and provide evidence that this mRNA species (RGS6LA3α1(+GGL)) encodes the RGS6B protein. Identification of this putative RGS6B isoform has the potential to open a new frontier of investigation into the mechanisms by which RGS6 contributes to both brain function as well as risk for neurological and neuropsychiatric disorders.

## Materials & Methods

### Mice

Generation of RGS6 knockout mice (RGS6^-/-^) has been described previously.^26^ Mouse brain tissues were isolated from adult (∼3 months of age), age-matched wild type and RGS6^-/-^mice that had been back-crossed onto a C57BL/6 background for 5 generations. Mice were euthanized by isoflurane inhalation followed by cervical dislocation and brains removed. Both male and female mice were used for tissue isolation as no sex differences in RGS6 protein expression have been noted. All animal procedures were carried out under a protocol approved by the University of Iowa Institutional Animal Care and Use Committee (IACUC).

### Western blotting

#### Mouse and Human Tissue Preparation

Mouse whole brain, frontal cortex, hippocampus, cerebellum, striatum, and/or ventral tegmental area (VTA) were rapidly dissected on ice and tissue samples snap frozen in liquid nitrogen for downstream processing. Tissue homogenates were prepared in RIPA buffer containing protease and phosphatase inhibitors (Millipore Sigma, St. Louis, MO, USA), quantified and probed as we previously described^7^. Twenty µg of protein per sample was subjected to SDS-PAGE and immunoblotting using standard techniques.

#### Cell Culture lysate preparation

At 24 h post-transfection, media was aspirated off cells and 100 µl of ice-cold RIPA buffer (150mM NaCl, 1% NP-40, 0.5% sodium deoxycholate, 0.1% sodium dodecyl sulphate, 50mM Tris-HCl: pH 8) containing protease (Roche, Basel, Switzerland) and phosphatase (Millipore Sigma) inhibitors was added to each well. Lysates were then placed in pre-chilled tubes, vortexed for 30 s, incubated on ice for 5 min, and centrifuged at 8,000 rpm (4⁰C) for 10 min. Supernatant was transferred to tubes containing 4x SDS-PAGE sample buffer and boiled for 5 min before loading onto gels.

#### Immunoblot visualization

Immunoblots were visualized using the Odyssey Imaging System with appropriate fluorescently labeled secondary antibodies (LI-COR Biosciences; Lincoln, NE, USA). Primary antibodies utilized included: rabbit anti-RGS6-L (in house generated polyclonal antibody against the RGS6L N-terminus),^25^ rabbit anti-RGS6-fl (in house generated polyclonal antibody against the full-length RGS6Lα1(+GGL) protein),^25^ rabbit anti-Gβ_5_ (a gift from William F. Simonds),^27^ rabbit anti-R7BP (a gift from Kirill A. Martemyanov), mouse anti-α-tubulin (CalBiochem, Cat# CP06), rabbit anti-β-Actin (Millipore Sigma, Cat# SAB5600204, RRID:AB_3097735), rabbit anti-GAPDH (Sigma, CAT# G9545). Densitometric quantification of western blots was performed utilizing Image J software (NIH) and expression of indicated proteins normalized to loading controls.

### Bioinformatic analyses

#### GWAS Catalog

GWAS reports implicating RGS protein family members in brain health were sourced from the GWAS catalog (https://www.ebi.ac.uk/gwas/). Studies were tabulated linking RGS protein genes to neurodegenerative disorders, neuropsychiatric disorders, neurovascular disease (e.g., stroke), cognitive function (e.g., education attainment, mathematical ability, impulse control), pain, sleep and/or circadian rhythms, brain structure (e.g., region volume), and or malignancy (e.g., glioblastoma, oligodendroglioma).

#### R7 family member expression in brain

To survey R7 family RGS protein gene expression across human brain regions, normalized RNA expression data (transcripts per million, TPM) for *RGS6*, *RGS7*, *RGS9*, *RGS11, GNB5,* and *RGS7BP* was extracted from the consensus human brain dataset of the Human Protein Atlas (https://www.proteinatlas.org/), which includes data from 13 major brain subregions. RGS protein expression in cell types of the frontal cortex was extracted from a single cell nuclear RNA seq (scRNAseq) cell atlas of the murine mouse brain available online at Dropviz.org.^28^ RGS6 protein expression in human cortex was extracted from the Broad Institute Single Cell Portal (https://singlecell.broadinstitute.org/) with raw data accessible in the Gene Expression Omnibus at GSE204684.

#### Phospho-prediction Analysis

Putative phosphorylation sites on RGS6LA3α1 (RGS6B) were identified using the Group-based Prediction System (GPS) 6.0 web-based application (https://gps.biocuckoo.cn/index.php) from Cuckoo Workgroup (Hubei, China)^29^. For our analysis, we inputted the unique C-terminal protein sequence present in RGS6LA3α1, the threshold was set to high, and predicted site/kinases were noted if their GPS prediction score was at least 0.8. A summary of hits is provided in **Figure S1** for the most frequently identified putative phosphorylation sites (S534, T535, S536) present in a consensus motif (RRRRSTS) at amino acids 530-536 in RGS6A3α1(+GGL).

#### Novel exon alignments

Evolutionary conservation of the novel RGS6 exons was assessed using UCSC Genome Browser (https://genome.ucsc.edu/) by aligning human RGS6 exon sequences to corresponding counterparts in other species.

### PCR amplification and identification of novel RGS6 cDNAs

Full-length cDNAs encoding novel RGS6 splice forms were amplified from a Marathon ready human brain cDNA library (Clontech; Mountain View, CA, USA) using a PCR-based strategy. In brief, we designed RGS6 specific primers, utilizing the NCBI Homo sapiens chromosome 14, GRCh38.p7 Primary Assembly (NC_000014.9), located in the RGS6L 5’UTR (RGS6L-5’) and in the 3’UTRs of terminal exons α and β (RGS6-αR2 and RGS6-βR3) (**Table S1**). RGS6 transcripts were amplified using Platinum Taq (ThermoFisher Scientific, Waltham, MA, USA) or EasyA Taq (TransGen Biotech, Beijing, China) and subsequently cloned into the pCR^TM^2.1-TOPO vector (ThermoFisher Scientific) using the manufacturer’s protocol. One-shot® Mach1^TM^-T1^R^ competent *E. coli* cells were transformed with the resulting RGS6-TOPO vectors and plated on Kanamycin (40 µg/µl) LB Agar plates. Transformants were screened for novel RGS6 transcripts by comparing, via gel electrophoresis, 1) the size of whole transcripts that were excised from the TOPO plasmid via restriction digest (HindIII and XbaI) to that of RGS6Lα1(+GGL) or 2) the size of partial transcripts amplified from isolated vectors using primers to exons 18 (RGS6-18) and β (RGS6-βR2) performing a PCR screen comparing generated transcripts to the predicted size of RGS6Lα1(+GGL) (**Table S1**). Once vectors containing novel transcripts were identified, they were sequenced, and their sequences analyzed for potential novel exons. Sanger DNA sequencing was performed at the Iowa Institute of Human Genetics, Genomics Division. Novel transcript sequences were confirmed as exons once the corresponding sequence had been identified in the human gene with the corresponding GT-AG splice donor-acceptor sites.

### PCR amplification of RGS6LA3(+GGL) transcript in mouse and human tissues

For these analyses RGS6LA3(+GGL) transcript was amplified from commercially available human cDNA libraries from muscle, kidney, heart, spleen, pancreas, and brain (Life Technologies, Waltham, MA) as well as from in house generated mouse tissue cDNA libraries. Mouse cDNA libraries were created using total RNA isolated from mouse tissues using the TRIzol® reagent (Life Technologies) following the manufacturer’s instructions. The quantity and quality of RNA were assessed using a NanoDrop 1000 (ThermoFisher Scientific). Total RNA was reverse-transcribed using Superscript™ II (Life Technologies). RGS6LA3 was amplified from all cDNA libraries using transcript specific primers directed toward the ATG start site in exon 2 (RGS6L-ATG-H and RGS6L-ATG-M) as well as the A3 exon (RGS6-A3R-H and RGS6-A3R-M) (**Table S1**).

### Preparation of pcDNA3.1 expression vector constructs

The novel RGS6LA3α1(+GGL) cDNA was PCR-amplified from the pCR^TM^2.1-TOPO vector using gene specific primers with restriction sites (RGS6L-ATG-H-KpnI and RGS6-αR2-H-NotI) to facilitate its cloning into pcDNA 3.1/*myc*-HisA (Life Technologies) (**Table S1**). The RGS6-αR2-NotI primer maintained the RGS6 TGA stop codon so RGS6 proteins are not *myc*-HisA-tagged. PCR products and vector then underwent restriction digestion and were ligated using the T4 DNA ligase system (New England Biolabs, Ipswich, MA, USA). A similar strategy was used to subclone RGS6Lα1(+GGL) and RGS6Lα2(+GGL) into the pcDNA 3.1/*myc*-HisA vectors, though a different 3’ primer was used to amplify these transcripts (RGS6-αR-NotI) (**Table S1**).

### Confirmation of RGS6LA3α1(+GGL) protein size via gel migration

HEK293T cells were plated in 12-well plates at 3.5 × 10 cell/well and cultured in high-glucose DMEM (Millipore Sigma) supplemented with 10% fetal bovine serum (Gibco, Waltham, MA, USA), 100 units/mL penicillin (Gibco), and 100 µg/mL streptomycin (Gibco). 24 h later, cells were transiently co-transfected using the Lipofectamine 2000 transfection reagent (ThermoFisher Scientific) with pcDNA 3.1/*myc*-HisA vectors (ThermoFisher Scientific) containing various RGS6 transcripts (total DNA = 1 µg/well). Culture media was exchanged for new supplemented DMEM media at 6 h following transfection. At 24 h post-transfection, cell lysates were prepared for Western blot analysis. Immunoblotting was performed utilizing an RGS6 antibody raised against the full-length RGS6Lα1 protein made in house.

### RGS6 isoform binding partner interactions and protein stability

HEK293T cells were plated and prepared for transfection as described above. 24 h later, cells were transiently co-transfected using the Lipofectamine 2000 system (ThermoFisher Scientific) with pcDNA^TM^3.1/*myc*-HisA (Life Technologies) vectors containing RGS6 transcripts, ± pcDNA3.1-HA vector containing Gβ_5_ (Gβ_5_-HA), and a pEGFP vector containing R7BP (R7BP-GFP; gifted from Dr. Kiril Martemyanov, Scripps, FL), or an empty pcDNA3.1 vector in a 2:1:1 ratio (total DNA = 1 µg/well). Transfection media was exchanged for new supplemented DMEM media at 6 h following transfection. At 24 h post-transfection cells were lysed and protein collected for immunoblotting. In a separate set of cells, media was removed at 24 h post-transfection and the cells were washed with warm 1 × DPBS. Serum free media containing 100 μM cycloheximide (Millipore Sigma) was subsequently added to each well. Following this at 0, 2, 6, 12, or 24h cells were lysed and protein collected for immunoblotting.

### RGS6 structure homology modeling

A structural model of RGS6LA3α1(+GGL) bound to Gβ_5_ was created using a previously described method^25^.

### Isolation of primary cortical astrocytes

Astrocytes were isolated from the whole cortex of neonatal (PN3-4) WT or RGS6^-/-^ mouse pups according to a previously published protocol^30^. Briefly, cortices were rapidly dissected from PN3-4 WT or RGS6^-/-^ mouse pups and placed into HBSS (Gibco) on ice. Cortices were then cut into small pieces and transferred to a 50 mL conical tube containing 10 mLs HBSS and 10 mLs 0.5% Trypsin (Gibco). Tubes were incubated for 30 mins at 37°C with shaking every 10 minutes. Next, dissociated cell suspensions were centrifuged at 300 × g for 5 min and the supernatant removed. Tissue was mechanically dissociated in 10 mL of astrocyte culture media (DMEM with high glucose + 10% FBS + Pennicillin/Streptomycin) by pipetting 20-30 times. Remaining tissue pieces were allowed to sediment and the resulting cell suspension transferred to poly-D-lysine (50 μg/mL) coated T75 flasks with an additional 10 mL of culture media. Cells were maintained at 37°C in a 5% CO_2_ incubator. After 2 days, the media was removed, cells rinsed with 1 X PBS, and media replaced. 1 week after isolation, the culture was enriched for astrocytes by shaking at 180 rpm for 30 mins at 37°C to remove microglia and then 240 rpm for 6 hours to remove oligodendrocytes. After ∼2 weeks or once cells reached confluency, they were re-plated for experiments.

### Generation of RGS6B-targed shRNA and miRNA and RGS6B knockdown in primary cortical astrocytes

We designed two shRNAis and Dr. Ryan Boudreau (University of Iowa) designed one miRNA selectively targeting the A3 exon found only in RGS6LA3α1(+GGL) using freeware. The unique selectivity of these RNAis for the A3 RNA sequence was confirmed by BLAST analysis. Complete synthesis of these shRNAis and miRNA in pFB-AAV-mU6mcs-CMV-eGFP-SV40pA was performed by GenScript (Piscataway, NJ, USA). Sequences are provided in **Table S2**.

### Glioma tumor lysates

Lysates from glioma tumor samples and control non-malignant brain tissues were obtained from Dr. Mahfoud Assem (University of Iowa, Department of Pharmaceutical Sciences and Experimental Therapeutics, Iowa City, IA). Tumor tissue samples were de-identified and assigned a numeric code to maintain patient confidentiality. Information regarding the tumor origin (astrocytic vs. oligodendrocytic) and stage were provided by the Department of Pathology (University of Iowa, Iowa City, IA) and are outlined in **Table S3**.

### MTT assay

Cell viability was measured by MTT reduction assay^17^. In brief, U87MG cells were seeded in 96-well plate at a density of 9,000 cells per well in 0.1 ml of phenol-free medium (Gibco). Transfection was performed 18 h post seeding with no DNA (blank), 100 ng of control DNA, or 100 ng of plasmids encoding RGS6L or RGS6B. At various times following transfection, 10 µl of 5 mg/ml of MTT (Thiazolyl Blue Tetrazolium Bromide, Biosynth T-3450) solution was added to the culture medium, and cells were incubated for 4 h at 37 °C. Formazan crystals in the viable cells were solubilized with 50 µl of dimethyl sulfoxide, and the absorbance at 570 nm was determined using a Cytation 5 plate reader (BioTek/Agilent). The survival rate is calculated using the following formula: Cell Viability (%)=100 × (ODsample−ODblank) / (ODcontrol−ODblank).

### Luciferase Assay

The ability of RGS6 isoforms to inhibit Gα_i_ was determined using a Nano-Glo Dual Luciferase Reporter Assay kit (Promega, Fitchburg, WI). In brief, astrocytes isolated from neonatal WT or RGS6^-/-^ mouse pups were seeded in 48-well plates that were coated with 50 µg/ml poly-D-lysine for 3 h at 37°C . Each well contained 45,000 cells in 0.25 ml of DMEM (Gibco) culture medium supplemented with 1 mM sodium pyruvate, 10% FBS, and 1x Pen/Strep. Transfection was performed 18 h post seeding using lipofectamine LTX Plus kit (Life Technologies) to deliver a total of 300 ng DNA. The DNA mixture contains 100 ng of a bicistronic reporter construct pNL[SV40-Fluc2, CRE-NlucP] and 200 ng of either control DNA, or plasmids encoding RGS6L or RGS6B. 48 h post-transfection, the culture medium was replaced with 0.2 ml of Opti-MEM (Gibco) containing either control vehicle (DMSO), or 100 nM isoproterenol hydrochloride (Millipore Sigma, I6504) with or without various concentrations of CCPA (MedChemExpress, Monmouth Junction, NJ, USA; HY-103185). 6 hours later, cells were lysed with 1x passive lysis buffer (Promega), followed by assaying both firefly (Fluc) and nano (Nluc) luciferases’ activities using a Nano-Glo Dual Luciferase Reporter Assay kit (Promega,). CRE promoter activity was expressed as relative nanoLuc (=Nluc activity/Fluc activity).

### Statistical Analysis

Data were analyzed by student’s t-test or one-way ANOVA or two-way ANOVA with post hoc adjustments to correct for multiple comparisons performed where appropriate. Normalized datasets were analyzed by one sample t-test evaluating deviation from a control group set to 1 or 100. Statistical analyses were performed using Prism software (GraphPad Software; La Jolla, CA, USA). Results were considered significantly different at *P*<0.05. Values are expressed as means ± S.E.M.

## Results

### The highly conserved RGS6B protein is expressed in several brain regions implicated in neurological disorders

In both mouse and human brain, multiple distinct RGS6 protein species are present that are lost in global RGS6^-/-^ mice (**Fig. 1C**). The 56 kDa band (magenta arrow) represents RGS6L(+GGL) isoforms, ubiquitously expressed throughout the body and identified in our initial cloning effort.^23,31^ However, in brain, multiple higher molecular weight bands can also be detected, that are larger than previously described RGS6 species. Most notable here, the 65 (orange arrow) and 69 kDa (green arrow) bands represent the phosphorylated and dephosphorylated brain specific RGS6, p-RGS6B and RGS6B, respectively.^31^ p-RGS6B, as with many phosphorylated proteins, is unstable. As human brain samples cannot be rapidly preserved, we typically do not see this band in human brain lysates or in murine lysates not immediately processed post-mortem. Furthermore, while RGS6L(+GGL) isoforms and RGS6B expression are conserved between mice and humans, it is important to note that the ratio of their expression is not. In human brain, RGS6B comprises a larger proportion of RGS6 protein species (**Fig. 1D**). In addition, the relative abundance of RGS6B vs RGS6L(+GGL) varies across mouse brain subregions with RGS6B being more prominent in the ventral tegmental area (VTA), striatum (STR), and frontal cortex (CTX) compared to the hippocampus (HIP) or cerebellum (CRB) (**Fig. 1E**). These data alluded to the possibility that RGS6B might have unique functional significance in select brain circuits and further supported our interest in determining its molecular identity.

### Molecular cloning of multiple novel RGS6 splice forms from human brain cDNA library

We hypothesized that RGS6B is the product of a novel RGS6 mRNA transcript, containing exons not identified in our initial cloning effort. In support of this hypothesis, northern blot analyses conducted in human brain and peripheral tissues show that there is at least one transcript predominantly expressed in the CNS that is larger than other previously identified RGS6 transcripts.^32^ Therefore, as a starting point in our investigation we performed a preliminary search of the NCBI database for transcripts that mapped to the *RGS6* gene but included potential novel exon sequences not identified in our initial cloning effort.^23^ One such sequence was uncovered (GenBank FLJ53896, AK303219.1), which contained three novel exons that mapped to the *RGS6* gene sequence between previously identified exon 18 and the α/β terminal exons (**Figs. 2A, 3**). We named the first two exons Alternate 1 and 2 (A1, A2) as they appeared to be optional exons which could be alternatively spliced in or out of the final RGS6 transcript. The third novel exon contained a stop codon and thus was termed θ in keeping with the nomenclature for previously identified terminal exons α-η.^23^

**Figure 2.**
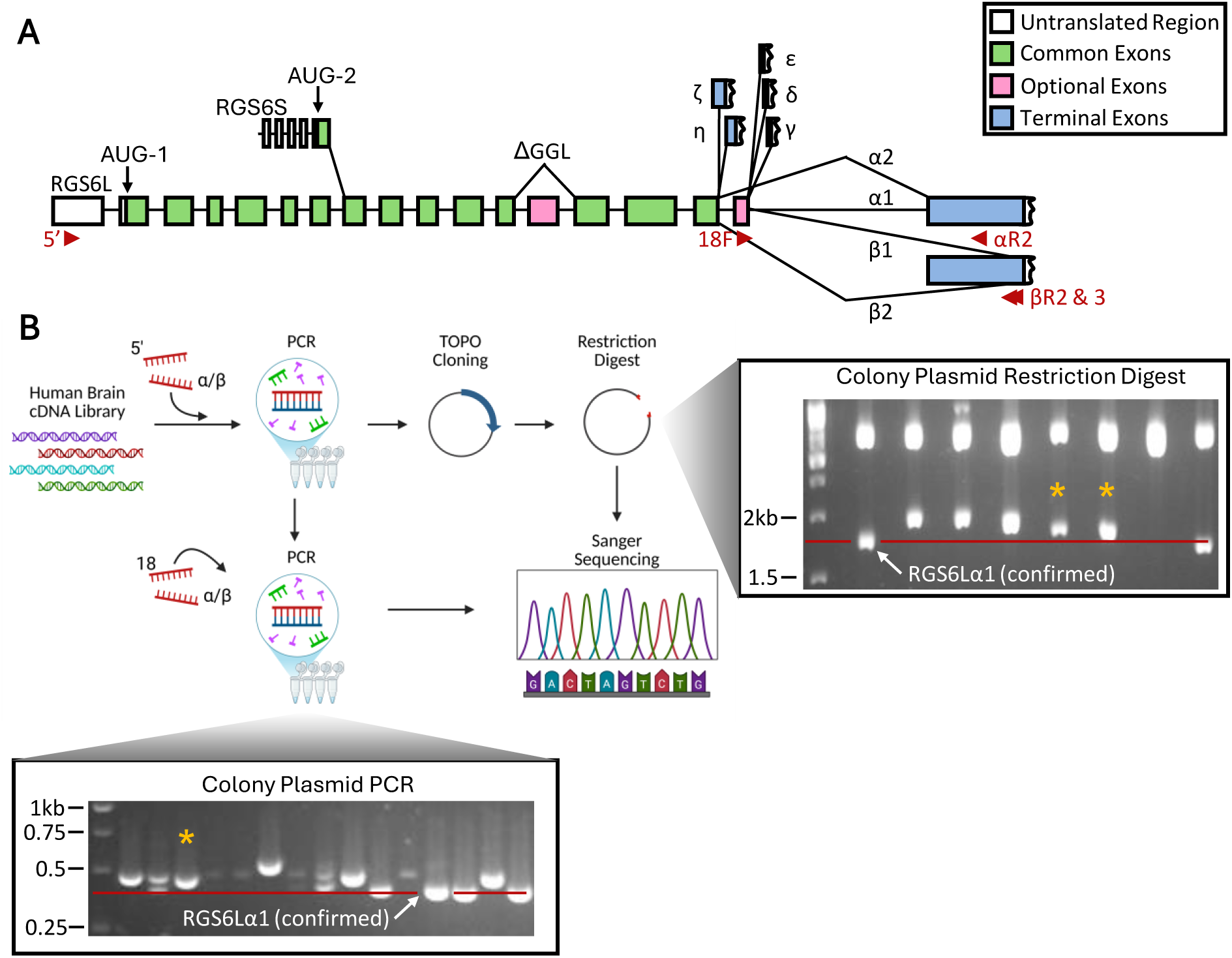
Molecular cloning of novel RGS6 splice forms from human brain. (**A**) Splicing diagram of known RGS6 splice forms. Location of primers used for PCR-based amplification of RGS6 transcripts from a human whole brain cDNA library are shown in red. (**B**) Representative plasmid restriction digest (right) or nested PCR (down) indicating positive hits whose size is larger than known RGS6 splice forms. Each gel has a confirmed RGS6Lα1(+GGL) transcript (largest known splice form) indicated as well as yellow asterisks denoting transcripts encoding RGS6LA3α1(+GGL) which encodes RGS6B.

**Figure 3.**
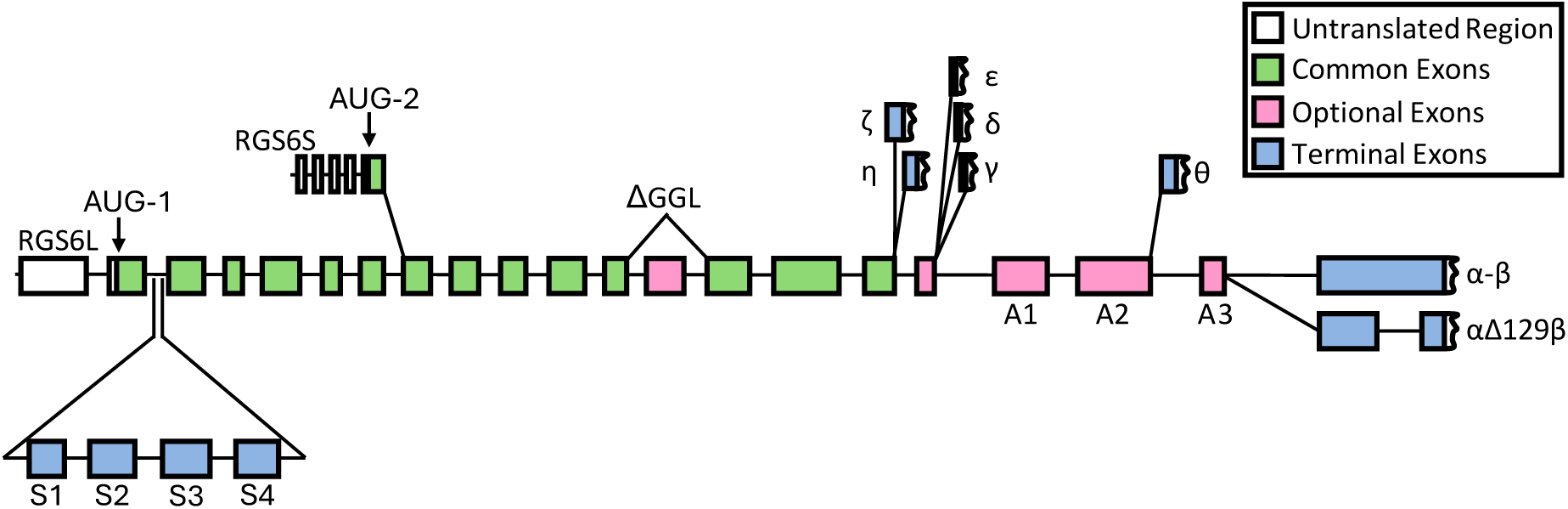
Updated RGS6 splicing diagram. Our recent cloning efforts have added 9 new exons to the RGS6 splicing scheme. Novel exons include: 4 small exons located between known exons 2 and 3 which each encode a premature stop codon (Stop (S)1-S4), 3 new “alternate” exons (A1-A3) located between known exons 18 and α/β , a new terminal exon (θ) located between A2 and A3, and finally an α/β terminal exon variant lacking a 129 bp central sequence (αΔ129β).

Together, these data supported our hypothesis that RGS6B may be the product of an RGS6 transcript containing novel exons. Therefore, we proceeded to a large scale RGS6 cloning project in which we sought to confirm the existence of the A1, A2, and θ exons, as well as identify other potential novel exon sequences that could contribute to the final RGS6B transcript. To this end, we designed a forward primer which recognized the 5’ UTR in exon 1 (RGS6-5’), present in all known RGS6L splice forms, as well as reverse primers to α or β terminal exons (RGS6-αR2 and -βR3), encoded by an overlapping sequence either containing (α) or lacking (β) a portion of the exon nucleotide sequence (**Fig. 2A**). These primer choices were made based on the following prior observations: 1) RGS6B contains protein sequence encoded by exons 1-6 and 2) the α/β terminal exon is the most 3’ sequence known to be included in RGS6 transcripts.^23,25^ PCR products amplified from the human brain cDNA library were subsequently cloned into TOPO vectors and transformed into bacteria for further amplification (**Fig. 2B**). Transformants were screened for novel RGS6 transcripts by comparing, via gel electrophoresis, 1) the size of whole transcripts enzymatically excised from the TOPO plasmid to that of RGS6Lα1(+GGL), the largest previously identified RGS6 transcript (**Fig 2B**, right), or 2) the size of partial transcripts, amplified from isolated vectors using primers to exons 18 (RGS6-18F) and β (RGS6-βR2), to that of RGS6Lα1(+GGL), whose 3’ end is composed of exon 18 spliced to the α/β terminal exon with no intervening sequence (**Fig 2B**, left).^23^

Using this approach, RGS6 transcripts containing 9 novel exons were identified. Through this screen we not only confirmed the existence of A1, A2, and θ which reside between known exons 18 and α/β but also discovered a fourth novel exon in the same region which we named A3, following our previously established nomenclature. In addition, 4 new exons were also identified between known exon 2 and exon 3 each of which, when included in the final transcript, encoded an early stop codon and hence were named Stop 1-4 (S1-S4). Finally, we identified a third splice variant for exon α/β that utilizes the same splice acceptor site needed to generate the β isoforms but an alternate donor site located within the α sequence. This results in an α/β exon hybrid sequence we termed αΔ129β (**Fig. 3**). Sequences for all novel exons have been included in **Table S4**. We also summarize in **Table S5** all RGS6 transcripts amplified, whether partial or complete, that contain these novel exon sequences. Though we highly doubt we have amplified and sequenced RGS6 transcripts encoding all possible exon combinations, our updated RGS6 mRNA splice diagram (**Fig. 3**), nonetheless, predicts a staggering number of possible RGS6 protein species with a conservative estimate, solely based on exon combinations we have confirmed, being over 400 possible sequences.

### RGS6B is encoded by the RGS6LA3α1(+GGL) transcript arising via novel exon inclusion

Though we now had evidence for numerous possible RGS6 protein isoforms, prior large-scale proteomic efforts have indicated that, in humans, most protein coding genes have a small number of dominant isoforms,^33^ an assertion consistent with the presence of only 3 RGS6L immunoreactive bands visible via western blot in human and mouse brain (**Fig. 1C**). Though it is possible that each band represents multiple protein species, we hypothesized that one or a small number of alternatively spliced mRNAs were likely responsible for producing the RGS6B protein band. Previously, we outlined five key attributes of the RGS6B protein that could aid in the identification of the transcript(s) encoding it.^25^ First, RGS6B is a larger protein than RGS6L, ∼69kDa. Second, RGS6B is expressed in both mouse and human brain tissue, indicating that the transcript encoding it must be derived from evolutionarily conserved regions within the *RGS6* gene. Third, RGS6B, like other RGS6L isoforms, is recognized by the RGS6-L antibody generated to the N-terminal amino acid sequence of RGS6L isoforms, indicating the RGS6B mRNA includes exon 2. Fourth, RGS6B is also recognized by the RGS6-18 antibody, generated to 14 amino acids encoded by exon 18, indicating that its transcript includes the variable exon 18 sequence. Fifth, RGSB contains a putative phosphorylation site that is not present in other RGS6L isoforms.

With the above-mentioned criteria in hand, we immediately excluded transcripts containing exons S1-S4 from our pool of possibilities as the proteins they were predicted to produce would be far smaller than 69 kDa. Furthermore, the S1-S4 exon sequences are poorly conserved (**Fig. S2A**) and are absent in mouse (data not shown). The A1, A2 and θ exons are also poorly conserved across vertebrate species (**Fig. S2B**) and, once again, largely absent in the mouse genome (data not shown). In contrast, the A3 exon is very highly conserved and, critically, nearly identical in mouse and human (**Fig. 4A, Fig. S2B**). Thus, we chose to focus on A3-containing transcripts.

**Figure 4.**
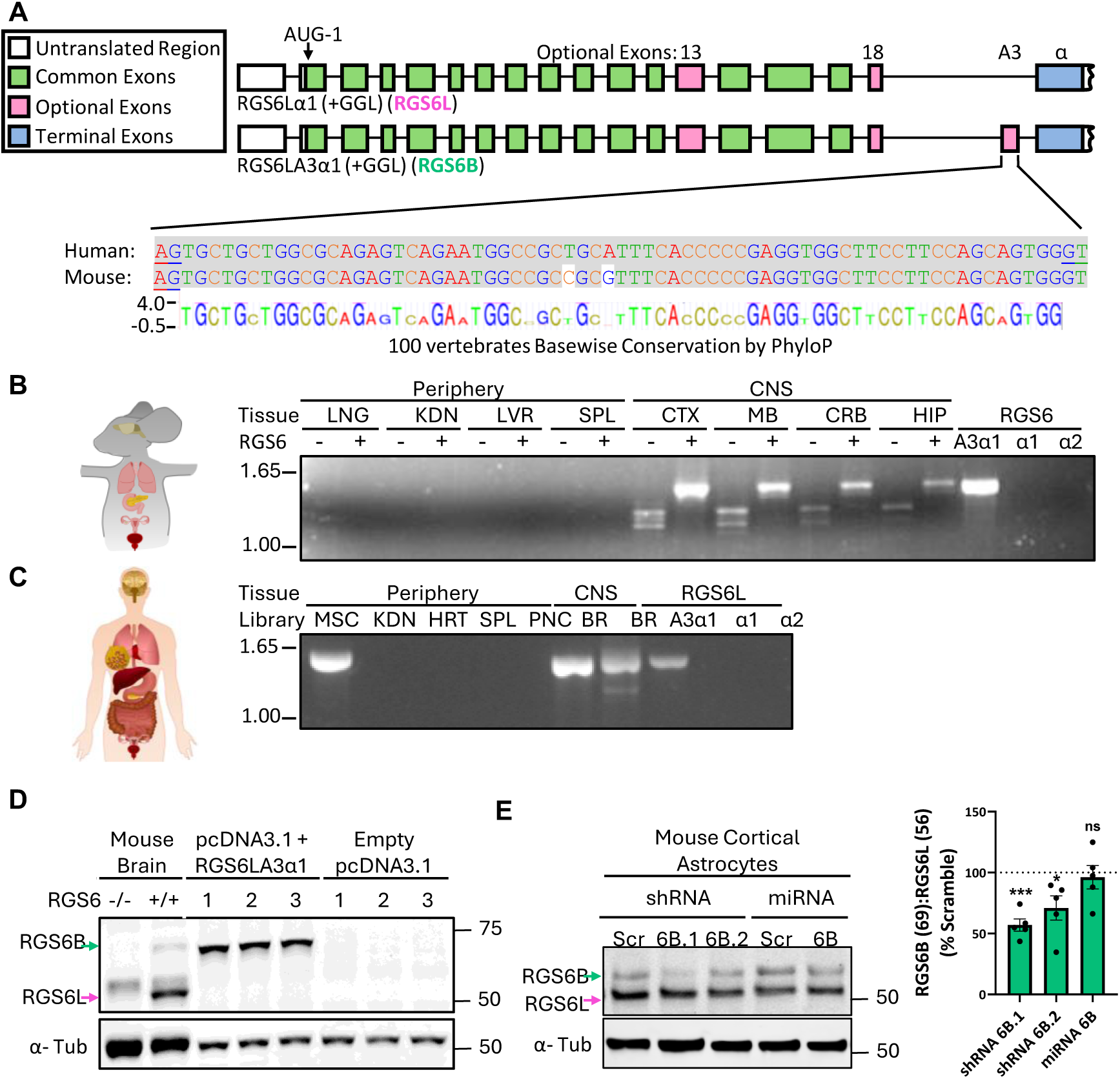
RGS6B is an RGS6L(+GGL) isoform containing A3 and the α terminal exon (RGS6LA3α1(+GGL)). (**A**) Schematic diagram of the exon splicing scheme comparing RGS6Lα1(+GGL) (RGS6L) and RGS6LA3α1(+GGL) (RGS6B). mRNA sequence conservation of exon A3 between mouse and human is depicted below with the consensus sequence from 100 vertebrate species extracted by PhyloP. PCR amplification of exon A3 containing mRNA transcripts from select mouse tissues (**B**) or human cDNA libraries (**C**). Plasmids encoding RGS6Lα1, RGS6Lα2 and RGS6LA3α1 (all +GGL) are used as negative and positive controls, respectively. LNG, lung; KDN, kidney; LVR, liver; SPL, spleen; CTX, cortex; MB, midbrain; CRB, cerebellum; HIP, hippocampus; MSC, muscle; HRT, heart; PNC, peripheral nervous system; BR, whole brain. (**D**) HEK293T cells were transfected with a plasmid encoding RGS6LA3α1(+GGL) to confirm co-migration with mouse RGS6B via immunoblotting. (**E**) shRNA or miRNA constructs targeting the A3 exon were introduced into mouse primary cortical astrocytes. The ratio of RGS6B: RGS6L was determined via immunoblot and quantified from 4 independent experiments. α Tubulin serves as a loading control for immunoblots. Data were analyzed by one sample t-test to detect deviation of each group from 100% (scramble RNAi control). *P<0.05, ***P<0.001. Data are expressed as mean ± S.E.M.

Through the course of our screen, we repeatedly identified a transcript (asterisks, **Fig. 2B**) which we named RGS6LA3α1(+GGL) to indicate it resembles the RGS6Lα1(+GGL) transcript identified in our initial cloning effort with the addition of novel exon A3. The RGS6LA3α1(+GGL) transcript matched an transcript published during our research in NCBI (NM_001370273) and met all the criteria we had previously established for RGS6B. First, RGS6LA3α1(+GGL) contains exons 1-6, which encodes the long N-terminus found in RGS6L isoforms. Second, the transcript contains exon 18. Third, the novel A3 exon is highly conserved and can be found in both mice and humans (**Fig. 4A**). Fourth, we found that RGS6LA3 transcript expression is largely restricted to CNS tissues in both mouse and human (**Fig. 4B** and **C**). The one exception being that RGS6LA3(+GGL) transcripts are also present in human skeletal muscle. However, the transcript may not be expressed at the protein level as we demonstrated that RGS6B is not expressed in mouse skeletal muscle.^25^ Fifth, HEK293T transfection experiments demonstrate that the RGS6LA3α1(+GGL) protein is comparable in size to RGS6B (**Fig. 4D**). Sixth, RGS6B has multiple putative phosphorylation sites absent in other RGS6 isoforms (described further in the next section, **Fig. S1**). Identifying an *in vitro* model to study RGS6B proved challenging as many cell lines lack endogenous RGS6 expression and primary cultures rapidly eliminate RGS6 due to its potent growth suppressive and pro-apoptotic actions.^17,34^ However, we do detect expression of RGS6L and RGS6B in primary neonatal mouse cortical astrocytes. Indeed, shRNA constructs targeting the A3 sequence caused RGS6B protein depletion from these cells while sparing RGS6L (**Fig. 4E**). These data provide definitive proof that the 69 kDa RGS6 immunoreactive band (RGS6B) is encoded by an A3-containing mRNA and, thus, represents RGS6LA3α1(+GGL).

### RGS6B protein contains a unique C-terminal extension

The only difference between RGS6Lα1(+GGL), abbreviated RGS6L, identified in our initial cloning effort^23^, and RGS6B, is the inclusion of the novel A3 exon. Inclusion of the A3 exon in the final transcript causes an alteration in the reading frame of terminal exon α resulting in an extension of the protein C-terminus (**Fig. 5A**). The novel C-terminus does not contain any known functional domains or similarity to other proteins and molecular modeling suggests it is a largely unstructured extension projecting out from the main body of the RGS6B protein in complex with Gβ_5_ (**Fig. 5B**). However, RGS6B’s novel C-terminus does contain multiple putative phosphorylation sites (bolded in **Fig. S1**) including a particularly strong hit (RRRRSTS) at amino acid residues 530-536 (underlined in **Fig. 5A**). Phospho-prediction analysis identified numerous kinases that could target these S/T residues (**Fig. S1**).

**Figure 5.**
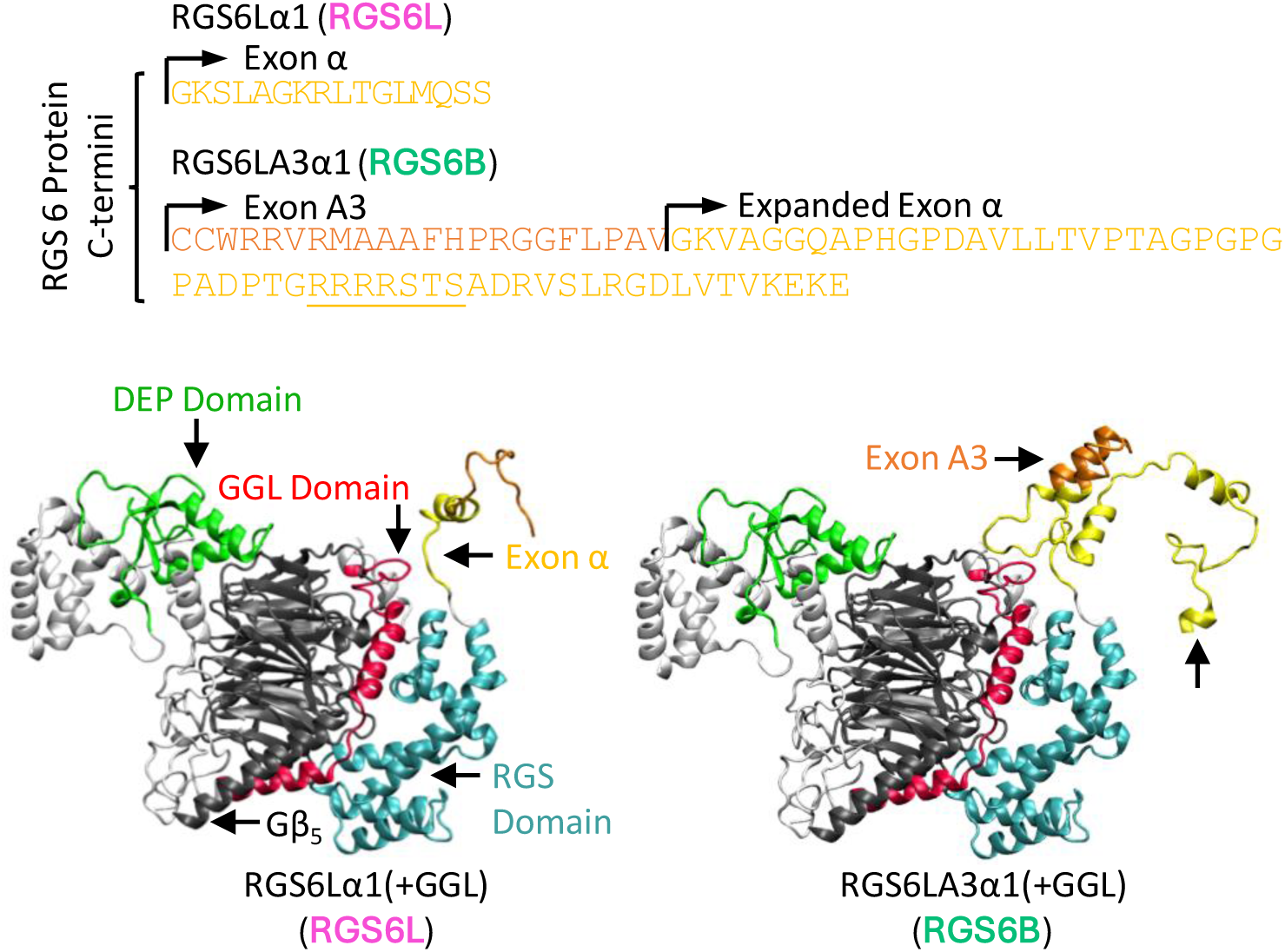
RGS6B contains a unique C-terminal extension. (**A**) Comparison of RGS6Lα1(+GGL) and RGS6LA3α1(+GGL) C-terminal sequences. The sequence encoded by exon A3 is highlighted in orange and the sequence encoded by terminal exon α in yellow. (**B**) Molecular modeling of RGS6Lα1(+GGL) (left) or RGS6LA3α1(+GGL) (right) in complex with Gβ5 (black). Domains are color coded (green = DEP, red = GGL, cyan = RGS). The structure generated from exon A3 is in orange and the structure generated from terminal exon α in yellow. The expanded C-terminus of RGS6LA3α1(+GGL) is largely unstructured and projects from the main body of the RGS6B protein adjacent to the RGS domain.

### RGS6B is more stable than other RGS6L(+GGL) isoforms and has an increased ability to stabilize RGS6 binding partner R7BP

As RGS6B contains intact DEP and GGL domains, we hypothesized that RG66B, like other RGS6L isoforms, would be able to interact normally with key RGS6 binding partners R7BP and Gβ_5_. To test this hypothesis, we transfected HEK293T cells with plasmids containing RGS6B or RGS6L alone or in combination with R7BP and/or Gβ_5_. As expected, due to their co-stabilizing properties, protein levels of RGS6, Gβ_5_, and R7BP were highest when all three proteins were co-expressed (**Fig. 6**). However, we noted two key differences between RGS6B and RGS6L when exogenously expressed. First, expression of RGS6B was consistently higher than that of RGS6L especially when both Gβ_5_ and R7BP were present. Second, whereas RGS6B and RGS6L were both equally capable of stabilizing Gβ_5_, RGS6B stabilized R7BP expression significantly more than RGS6L (**Fig. 6**).

**Figure 6.**
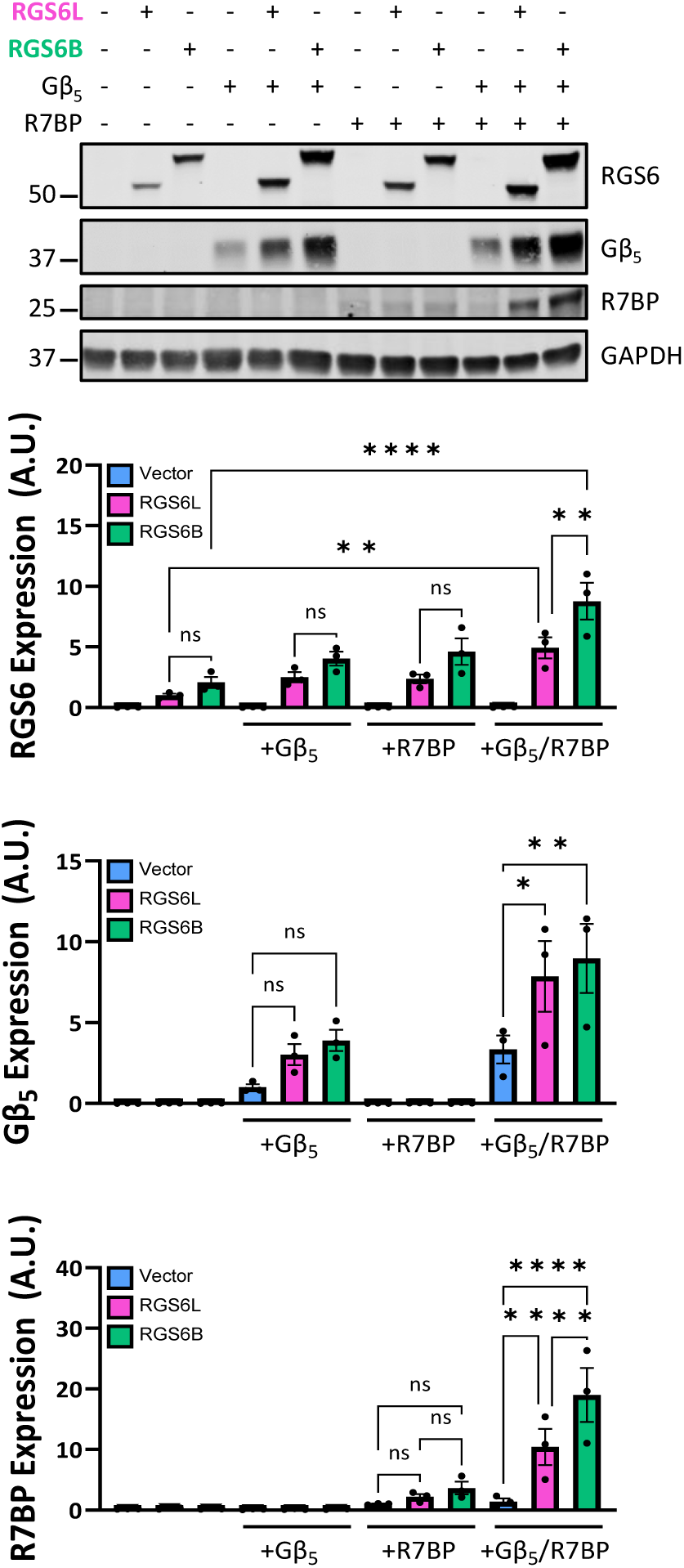
Both RGS6B and RGS6Lα1(GGL) have co-stabilizing functional interactions with Gβ5 and R7BP, binding partners for R7 family RGS proteins. Plasmids encoding RGS6B (RGS6LA3α1(+GGL)) or RGS6Lα1(+GGL) were co-transfected into HEK293T cells ± R7BP, Gβ5 or both. Protein expression was determined via immunoblot with GAPDH serving as the loading control. Densitometric quantification from 3 independent experiments is shown. Data were analyzed by one-way ANOVA with Sidak’s post-hoc test.*P<0.05, **P<0.01, ****P<0.0001. ns = not significant. Data are expressed as mean ± S.E.M.

As the same promoter is driving expression of both RGS6B and RGS6L when expressed exogenously, we hypothesized that the difference in RGS6 expression level was due to differential protein stability. This hypothesis was confirmed using cycloheximide chase experiments (**Fig. 7**). RGS6B displays a longer protein half-life as compared to RGS6L especially in the absence of Gβ_5_ and R7BP (**Fig. 7A-B**). Indeed, after 24 hours of cycloheximide treatment, RGS6B protein expression is largely maintained while little to no RGS6L is detectable (**Fig. 7C**).

**Figure 7.**
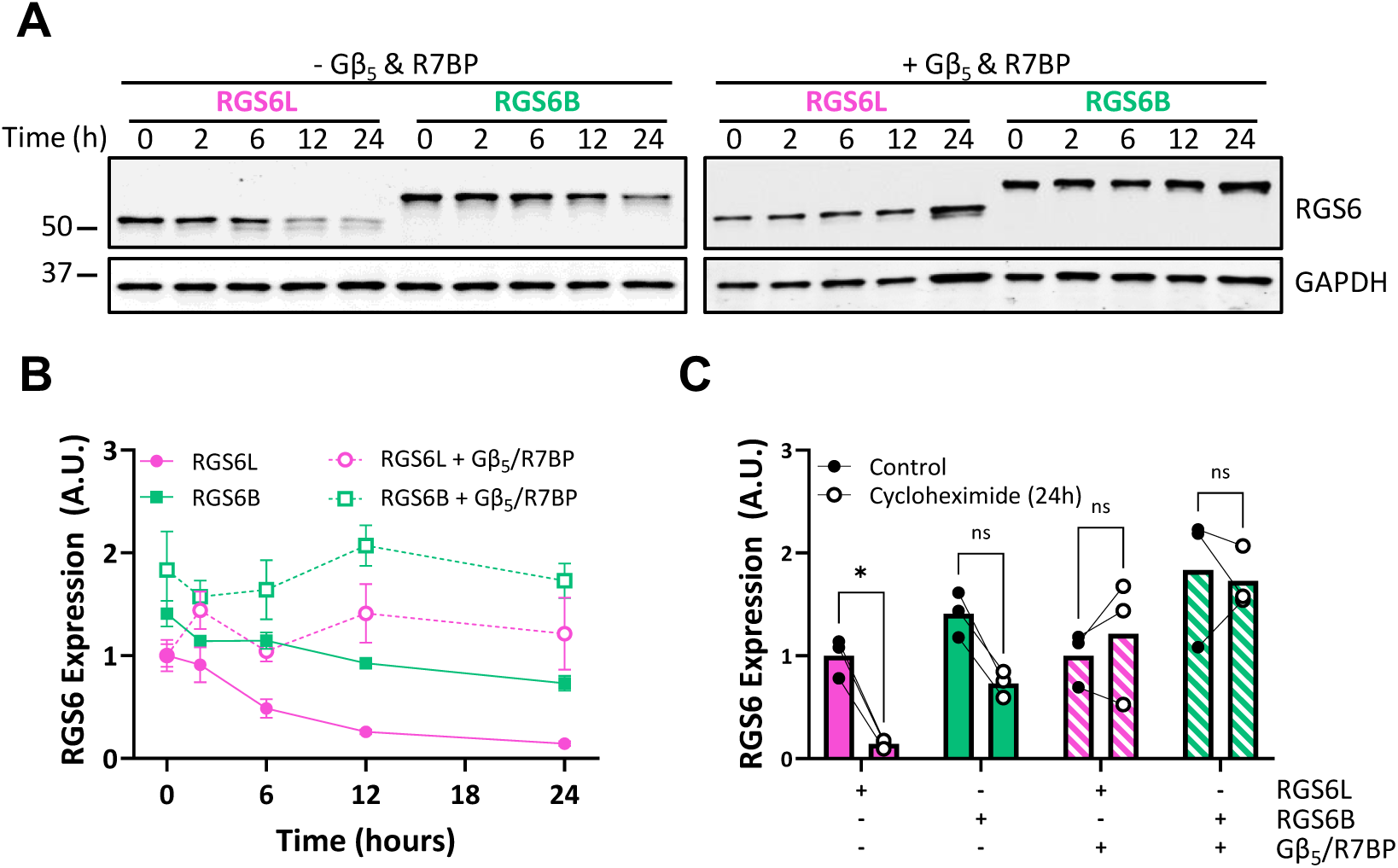
RGS6B is more stable than RGS6Lα1(+GGL). (**A**) HEK293T cells were transfected with plasmids encoding RGS6B (RGS6LA3α1(+GGL)) or RGS6L (RGS6Lα1(+GGL)) ± R7BP/Gβ5. 24 hours later, cells were treated with cycloheximide (100 μM) and lysed at 0, 2, 6, 12, or 24 hours following inhibition of translation. Expression of RGS6 was determined via immunoblotting with GAPDH serving as the loading control. RGS6 expression was quantified via densitometry and data are provided as (**B**) full time course or (**C**) control vs 24-hours post-treatment (n=3). Data were analyzed by two-way ANOVA with Sidak’s post-hoc test. .*P<0.05. ns = not significant. Data are expressed as mean ± S.E.M.

### RGS6B maintains non-canonical cytotoxic action but lacks the capacity for G protein regulation

Next, we wished to evaluate possible functional divergence between RGS6L and RGS6B isoforms. RGS6L functions as a tumor suppressor due to its potent G protein-independent pro-apoptotic and growth suppressive actions.^17,21,22^ ^34^ In the cortex of mouse (**Fig. 8A**) and human (**Fig. 8B**) brain, RGS6 mRNA is detectable in multiple cell types including pyramidal neurons and interneurons, but also glia with highest glial expression in astrocytes. In fact, RGS6 is one of the most prevalent RGS proteins in cortical astrocytes (**Fig. S3**). As we have previously observed in breast^21^ and bladder^22^, RGS6 protein is depleted in tumor samples from human patients with malignant gliomas of both astrocytic and oligodendrocytic origin relative to healthy controls (**Fig. 8C, 8D**). Notably, loss of both RGS6L and RGS6B protein species was observed in gliomas, and we noted several instances wherein the migration of RGS6B shifted relative to control samples, which could indicate mutation, posttranslational modification, or an alteration in mRNA splicing (**Fig. 8C, green asterisks**). This phenomenon was frequently observed in high grade glioblastomas (6 of 8) (**Fig. 8C**; **Table S3**). Consistent with their loss in cancer, both RGS6L and RGS6B impaired the survival of U87MG cells indicating that RGS6B, like RGS6L, acts as a growth suppressor (**Fig. 8E**).

**Figure 8.**
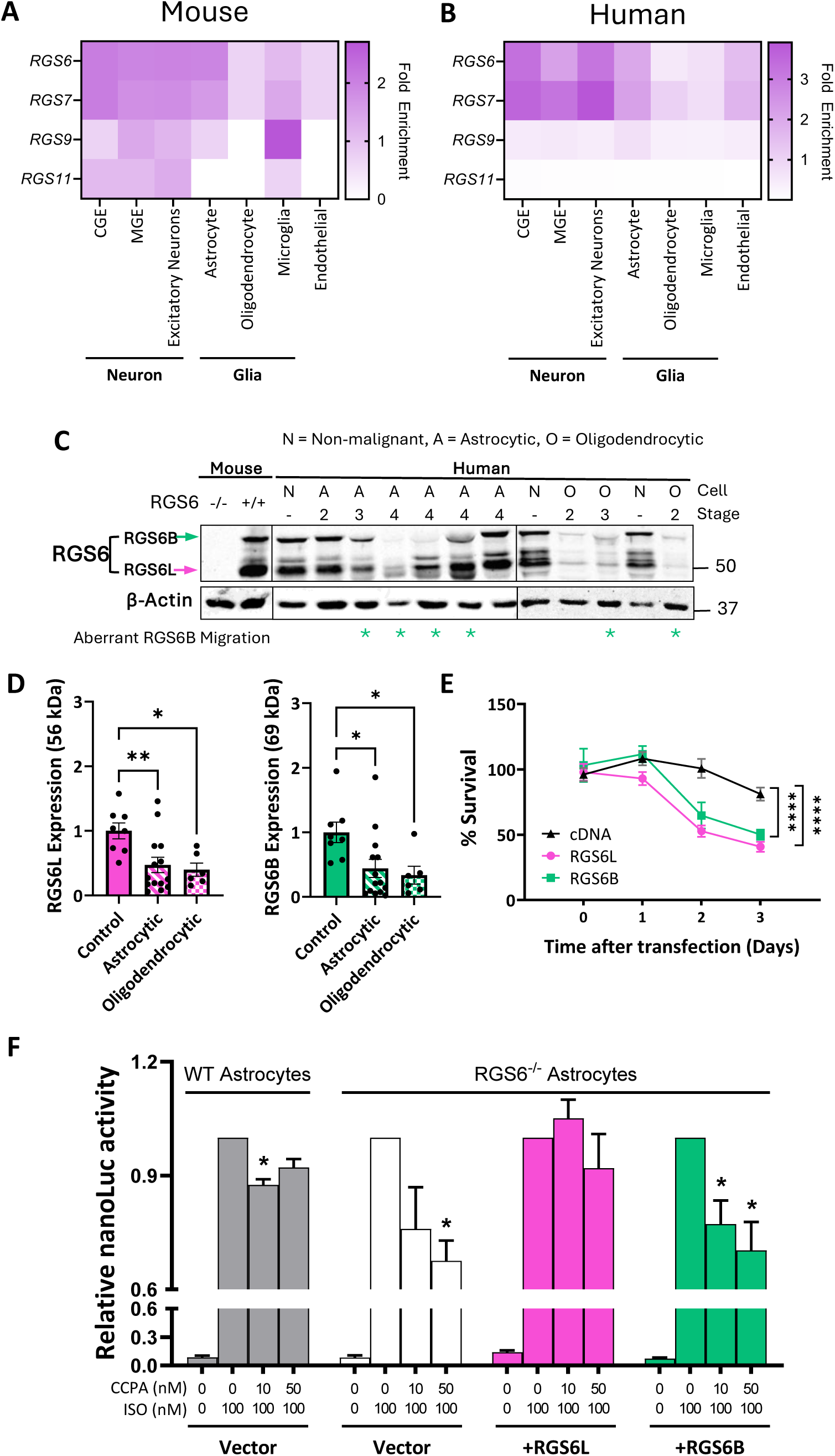
RGS6B suppresses growth of cancer cells but fails to impact Gαi/o signaling. mRNA expression of R7 family RGS proteins across cell types in the frontal cortex in (**A**) mouse [Dropviz.org] and (**B**) human [https://singlecell.broadinstitute.org/]. (**C**) Immunoblotting and (**D**) densitometric quantification of RGS6L (56 kDa) and RGS6B (69 kDa) expression in malignant tumors of astrocytic (n=14) or oligodendrocytic (n=6) origin isolated from human cancer patients relative to non-malignant controls (n=8). β-Actin serves as a loading control. Green stars indicate lanes where aberrant migration of the RGS6B band was noted. (**E**) Survival of U87MG cells at various times following RGS6L, RGS6B or control DNA transfection as measured by MTT assay (n=4). (**F**) RGS6-/- or WT murine cortical astrocytes were co-transfected with a dual luciferase CREB reporter construct ± RGS6L, RGS6B, or vector DNA. 48 hours later, cells were then treated with vehicle (DMSO) or a cocktail of isoproterenol (100 nM) with varying concentrations of the A1R agonist CCPA (10-50 nM). Luciferase was used to evaluate the ability of RGS6 isoforms to inhibit CCPA-induced blockade of CRE promoter activity in the luciferase reporter construct. Data were analyzed by one-way (**C**) or two-way (**E**) ANOVA with Dunnett’s or Sidak’s post-hoc test, respectively. Data in (**F**) were analyzed by one sample t-test assessing deviation from 1. *P<0.05, **P<0.01, ****P<0.0001. ns = not significant. Data are expressed as mean ± S.E.M.

Our work, and that of others, has demonstrated a key role for RGS6 in regulation of Gα_i/o_-coupled GPCRs in the nervous system via its canonical function as a GTPase activating protein (GAP).^7,9,14^ To test RGS6B’s G protein regulatory capacity, we turned again to the astrocyte as murine cortical astrocyte cultures express both RGS6L and RGS6B protein (**Fig. 4E**). We monitored cAMP response element binding protein (CREB)-dependent transcriptional activation of a luciferase reporter in the presence of isoproterenol to stimulate Gα_s_-coupled β1 adrenergic receptors and increasing concentrations of 2-chloro-N6-cyclopentyladenosine (CCPA), a selective agonist for Gα_i/o_-coupled adenosine A1 receptors (A1Rs). A1R-driven inhibition of reporter activity was absent in RGS6^-/-^ astrocytes following reintroduction of RGS6L (**Fig. 8F**). However, CCPA did impair cAMP signaling in RGS6 null astrocytes transfected with RGS6B (**Fig. 8F**). In fact, we noted that co-transfection of RGS6B with RGS6L resulted in attenuation of RGS6’s GAP activity indicating that RGS6B can function as a dominant negative in this system (**Fig. S4**). This observation is consistent with the minimal impact of CCPA on reporter activity in WT cells, which express both RGS6L and RGS6B (**Fig. 8F, 4E**).

## Discussion

The central aim of this study was to determine the molecular identity of a novel, highly conserved RGS6 protein isoform exclusively detected in the nervous system (RGS6B).^25^ This work describes the cloning and initial functional characterization of a new RGS6 transcript which encodes RGS6B. This transcript was named RGS6LA3α1(+GGL) to indicate it most closely resembles the previously identified RGS6Lα1(+GGL) transcript.^23^ The only difference between the two transcripts is the inclusion of the novel, highly conserved, A3 exon. A3 inclusion in the final transcript is ultimately predicted to cause a shift in the reading frame of terminal exon α resulting in an unstructured extension of the protein C-terminus.

### Though the RGS6 gene is predicted to encode a staggering number RGS6 protein isoforms, only a handful, which largely differ in their C-termini, appear to be expressed

Through a second large scale RGS6 cloning project, described here, we uncovered RGS6 transcripts containing 9 novel exons (S1-4, A1-3, θ, and αΔ129β) not identified in our initial cloning effort.^23^ While we highly doubt we have amplified and sequenced RGS6 transcripts encoding all possible exon combinations, our updated RGS6 mRNA splice diagram (**Fig. 3**), nonetheless, predicts 420 possible RGS6 splice forms. Whereas we have evidence for numerous possible RGS6 protein isoforms, prior large-scale proteomic efforts have indicated that, in humans, most protein coding genes have a small number of dominant isoforms.^33^ Consistent with this assertion, our previous comprehensive analysis of RGS6 protein expression in mice demonstrated that, of the four possible RGS6 protein classes identified previously, RGS6L(+/- GGL) and RGS6S(+/-GGL), only the RGS6L(+GGL) class is detected *in vivo*. Furthermore, prior work and the data described herein indicate that the novel brain-specific 65 and 69kDa RGS6 protein bands identified via western blot represent the same RGS6B protein, in phosphorylated and dephosphorylated states, respectively, encoded by a single mRNA splice form, RGS6LA3α1(+GGL).^25^

It is important to note that in both of our cloning efforts we have shown that the greatest diversity in RGS6 pre-mRNA alternate splicing occurs at the 3’ end of the transcript (**Fig 3A** and **4**). In fact, when looking at predominantly expressed RGS6L(+GGL) transcripts they differ only in their terminal exon sequences. These terminal exons in turn encode relatively short protein sequences, 4-34 amino acids in length with some overlap, making it difficult to distinguish between the RGS6L(+GGL) isoforms. Very little is known regarding the impact of the subsequent variable C-terminal sequences on overall RGS6 protein function. Given our current evidence for an even more diverse array of exon incorporation(s) in the 3’ end of the RGS6 transcript, as well as our evidence that RGS6B differs from other RGS6L(+GGL) proteins only in its C-terminus, it becomes even more important to understand the impact of these C-terminal sequences on RGS6 activity, expression, localization and effector binding. As RGS6B appears to lack functional G protein-regulating activity (**Fig. 8F**), it stands to reason that the unique C-terminus of RGS6LA3α1 precludes interaction with Gα_i/o_ or otherwise impairs RGS domain activity. Notably, the majority of novel 3’ exons (A1, A2, and θ) identified in this study are not well conserved and are largely present in humans and higher primates. Only the A3 exon showed wide-spread conservation. Therefore, there is also a question of what function, if any, the A1, A2, and θ exons confer selectively to RGS6 protein function in higher mammals, given the RGS6 protein species detectable via immunoblotting are the same in mouse and humans.

### RGS6B’s novel C-terminal extension promotes protein stability, enhances R7BP functional interaction, and impairs its ability to negatively regulate Gi/o

RGS6B’s novel C-terminal extension alters protein function compared to the other RGS6L(+GGL) isoforms (**Figs. 6, 7, and 8E**). There is precedence for RGS6 splice forms differing in their function. Namely, those RGS6 splice forms missing exon 13 (ΔGGL, **Fig. 3A**) lack an intact GGL domain and are unable to bind Gβ_5_.^6,23^ As such, compared to RGS6L(+GGL) isoforms, RGS6L(-GGL) isoforms are less stable and are not detectable *in vivo*.^25^ Furthermore, RGS6S isoforms predominantly exhibit nuclear or nucleolar localization unlike RGS6L isoforms, which remain in the cytoplasm in the absence of Gβ ^23^. Similarly, other RGS pre-mRNA (RGS3, 9 and 12) are also prone to alternative splicing, producing multiple protein isoforms that differ in their sub-cellular localization, protein-protein interactions and cellular signaling^35–40^. With this data in mind, it is not surprising to find that RGS6B functions do not completely overlap with that of the other RGS6L(+GGL) isoforms.

RGS6B is more stable, its half-life is significantly longer than that observed for other RGS6L(+GGL) isoforms (**Fig. 7**). Mechanism(s) controlling RGS6 intracellular turnover remain largely unexplored, and, as a result, it is difficult to speculate as to how RGS6B’s extended C-terminus might influence protein half-life. This difference may be, at least in part, due to RGS6B’s differential ability to stabilize R7BP (**Fig. 6**). Indeed, knockout of R7BP has been shown to cause RGS9-2 depletion, a phenotype attributed to unmasking of a motif facilitating constitutive destruction of RGS9-2 by lysosomal cysteine proteases.^41,42^ Furthermore, the domain mediating RGS9-2 degradation and/or R7BP-dependent stabilization is categorized as “unstructured” and located at the C-terminus of the protein where heat shock cognate protein 70 (Hsc70) binds,^43^ observations that parallel unique features of RGS6B and suggest a link between the C-terminal RGS6B domain, R7BP binding, and protein degradation.

R7BP plays a key role in recruiting R7 family members to the plasma membrane where it facilitates their ability to regulate GPCRs and G protein effectors such as G protein activated inwardly rectifying potassium (GIRK) channels.^44–47^ Thus, enhanced R7BP binding would be expected to enhance RGS6B-dependent GPCR inhibition. However, our data demonstrate that RGS6B, unlike RGS6L, is unable to block Gα_i/o_ signaling and acts in a dominant negative fashion to prevent RGS6L from acting as a GAP in astrocytes (**Figs. 8E, S4**). Thus, instead, it is possible that RGS6B acts as a sink for endogenous R7BP interfering with R7BP-dependent facilitation of GPCR regulation by RGS6L or, alternatively, the RGS6B-R7BP complex may have G protein-independent effectors at the plasma membrane.

Thus, RGS6B, which itself lacks the ability to negatively regulate Gi/o signaling, can function to promote Gi/o signaling in cells, such as astrocytes and neurons, that express RGS6L via its dominant negative actions. Though RGS6 proteins were discovered as proteins that function to attenuate G protein signaling, RGS6B is the first example of an RGS protein member capable of enhancing G protein signaling. The inclusion of the novel C-terminal domain of RGS6B, while not affecting its ability to modulate G protein-independent cytotoxicity, presumably is responsible for the loss of its G protein-regulating activity and its dominant negative activity towards RGS6L. The present observations suggest that increases in the expression of RGS6B vs other RGS6 isoforms can flip the role of RGS6 proteins from inhibiting to stimulating Gi/o signaling. Given the critical role of Gi/o signaling in the central nervous system, and the robust expression of RGS6L and RGS6B in neurons and astrocytes, this would be expected to have significant effects on Gi/o-regulated neuronal functions, including the release and re-uptake of neurotransmitters and overall neuronal excitability. Future studies will seek to investigate the structural basis underlying these actions of RGS6B.

### Putative phosphorylation sites in RGS6B’s novel C-terminal extension hint at potential its potential role in neurological disorders

We previously demonstrated that RGS6B, unlike other RGS6L(+GGL) protein species, exists in phosphorylated and dephosphorylated states^25^. Specifically, RGS6B can undergo Ca^2+^-dependent phosphoregulation resulting in a detectable mobility shift during SDS-PAGE (69 → 65 kDa).^25^ Importantly, exon A3, when spliced into the RGS6Lα1(+GGL) transcript, causes an alteration in the reading frame of terminal exon α producing RGS6B’s unique C-terminal extension (**Fig. 6A**), which contains multiple novel putative phosphorylation sites (**Fig. S1**) including a particularly strong hit (RRRRSTS) at amino acid residues 530-536 (underlined in **Fig. 6A**). Phospho-prediction analysis has identified numerous kinases that could target these S/T residues (**Fig. S1**). This analysis revealed kinases that act as oncogenes (e.g., AKT1) or tumor suppressors (e.g., CHK1/2) consistent with the loss of RGS6B in glioma and its ability to suppress growth of U87MG glioma cells. Several kinases were also identified which have been previously associated with nervous system function and dysfunction, including: Ca^2+^/calmodulin-dependent protein kinases (CAMK) IIα and IV as well as Leucine-rich repeat kinase 2 (LRRK2). CAMK’s II and IV have been implicated in autism spectrum disorder (ASD) and are known regulators of neurodevelopment and synaptic plasticity (as reviewed by Kaiser et al.^48^). RGS6 modulates dendrite outgrowth and neuronal differentiation through its interaction with stathmin-like protein 2 (SCG10).^49^ Although these experiments utilized RGS6L isoforms, it is possible that RGS6B might be a CaMK target linking Ca^2+^ oscillations to dendritic remodeling. LRRK2, a monogenic risk factor for Parkinson’s Disease (PD),^50^ was also identified as a putative RGS6B kinase. As knockout of all RGS6 isoforms triggers the age-dependent degeneration of dopaminergic neurons characteristic of PD that is accompanied by LRRK2 up-regulation,^8,51^ a functional interaction between RGS6 and LRRK2 in an isoform-specific manner is an intriguing potential avenue for further research, especially given that the midbrain is a site of enriched RGS6B expression.

## Conclusions

Greek philosopher Aristotle said, “nature does nothing in vain”. We now estimate that there are 420 possible RGS6 splice forms, an astounding diversity unprecedented in the RGS protein family. Though not a record-breaking number when considering the entirety of the human genome, this inherent complexity in RGS6 protein generation must serve a purpose especially given many of the novel RGS6 splice forms we identified here are exclusively present in the nervous system. As *RGS6* has been heavily implicated in brain disorders in humans (**Fig. 1A**) and studies of global RGS6^-/-^ mice have repeatedly confirmed the highly pleiotropic nature of RGS6 biology^52,53^, identifying both those splice forms most favored across cell types as well as the unique functionality of RGS6 protein isoforms will be critical for establishing a comprehensive understanding of the role of RGS6 in nervous system function and dysfunction. Indeed, we recently demonstrated that splice acceptor site variant in the RGS6 gene, which results in congenital cataracts, intellectual disability, and microcephaly^24^, favors production of a subset of RGS6 splice forms^31^ providing key evidence linking RGS6 pre-mRNA splicing to disease pathogenesis. Though it has been known for over 20 years that the greatest diversity of alternate splicing events involved in processing RGS6 pre-mRNA happen at the 3’ end, very little is known regarding the impact of the C-terminal variant sequences on RGS6 protein function. Given our evidence for an even more diverse array of exon incorporation(s) in this part of the protein, efforts to understand the impact of these sequences on RGS6 activity, expression, localization and effector binding might provide valuable insight into the physiological and pathophysiological functions of RGS6. These studies are now possible given our ability to selectively deplete RGS6B in brain regions using AAV expressing RGS6B RNAis or overexpress RGS6B (in RGS6^-/-^ mice) in discrete brain regions.

## Acknowledgements

The work presented in this manuscript was supported by NIH AA025919 (RAF) and a Dee Silver Pilot Award in Neurodegeneration (AS).

## References

1. Cross-Disorder Group of the Psychiatric Genomics Consortium. Electronic address, p.m.h.e. & Cross-Disorder Group of the Psychiatric Genomics, C. Genomic Relationships, Novel Loci, and Pleiotropic Mechanisms across Eight Psychiatric Disorders. Cell 179, 1469–1482 e1411 (2019).

2. Cerezo, M., et al. The NHGRI-EBI GWAS Catalog: standards for reusability, sustainability and diversity. Nucleic Acids Res 53, D998–D1005 (2025).

3. Martemyanov, K.A., Yoo, P.J., Skiba, N.P. & Arshavsky, V.Y. R7BP, a novel neuronal protein interacting with RGS proteins of the R7 family. J Biol Chem 280, 5133–5136 (2005).

4. Drenan, R.M., et al. R7BP augments the function of RGS7*Gbeta5 complexes by a plasma membrane-targeting mechanism. J Biol Chem 281, 28222–28231 (2006).

5. Nini, L., et al. R7-binding protein targets the G protein beta 5/R7-regulator of G protein signaling complex to lipid rafts in neuronal cells and brain. BMC Biochem 8, 18 (2007).

6. Chen, C.K., et al. Instability of GGL domain-containing RGS proteins in mice lacking the G protein beta-subunit Gbeta5. Proc Natl Acad Sci U S A 100, 6604–6609 (2003).

7. Maity, B., et al. Regulator of G protein signaling 6 (RGS6) protein ensures coordination of motor movement by modulating GABAB receptor signaling. J Biol Chem 287, 4972–4981 (2012).

8. Luo, Z., et al. Age-dependent nigral dopaminergic neurodegeneration and alpha-synuclein accumulation in RGS6-deficient mice. JCI Insight 5(2019).

9. Stewart, A., et al. Regulator of G-protein signaling 6 (RGS6) promotes anxiety and depression by attenuating serotonin-mediated activation of the 5-HT(1A) receptor-adenylyl cyclase axis. FASEB J 28, 1735–1744 (2014).

10. Spicer, M.M., et al. Regulator of G protein signaling 6 mediates exercise-induced recovery of hippocampal neurogenesis, learning, and memory in a mouse model of Alzheimer’s disease. Neural Regen Res 20, 2969–2981 (2025).

11. Stewart, A., et al. Regulator of G protein signaling 6 is a critical mediator of both reward-related behavioral and pathological responses to alcohol. Proc Natl Acad Sci U S A 112, E786–795 (2015).

12. Spicer, M.M., et al. Regulator of G protein signaling 6 (RGS6) in dopamine neurons promotes EtOH seeking, behavioral reward, and susceptibility to relapse. Psychopharmacology (Berl) 241, 2255–2269 (2024).

13. Luo, H., et al. Receptor-dependent influence of R7 RGS proteins on neuronal GIRK channel signaling dynamics. Progress in neurobiology 243, 102686 (2024).

14. DeBaker, M.C., et al. RGS6 negatively regulates inhibitory G protein signaling in dopamine neurons and positively regulates binge-like alcohol consumption in mice. Br J Pharmacol 180, 2140–2155 (2023).

15. Yang, J., et al. G-protein inactivator RGS6 mediates myocardial cell apoptosis and cardiomyopathy caused by doxorubicin. Cancer Res 73, 1662–1667 (2013).

16. Huang, J., Yang, J., Maity, B., Mayuzumi, D. & Fisher, R.A. Regulator of G protein signaling 6 mediates doxorubicin-induced ATM and p53 activation by a reactive oxygen species-dependent mechanism. Cancer Research 71, 6310–6319 (2011).

17. Maity, B., et al. Regulator of G protein signaling 6 (RGS6) induces apoptosis via a mitochondrial-dependent pathway not involving its GTPase-activating protein activity. J Biol Chem 286, 1409–1419 (2011).

18. Mahata, T., et al. Hepatic Regulator of G Protein Signaling 6 (RGS6) drives non-alcoholic fatty liver disease by promoting oxidative stress and ATM-dependent cell death. Redox Biol 46, 102105 (2021).

19. Sengar, A.S., et al. RGS6 drives cardiomyocyte death following nucleolar stress by suppressing Nucleolin/miRNA-21. J Transl Med 22, 204 (2024).

20. Huang, J., et al. RGS6 suppresses Ras-induced cellular transformation by facilitating Tip60-mediated Dnmt1 degradation and promoting apoptosis. Oncogene 33, 3604–3611 (2014).

21. Maity, B., et al. Regulator of G protein signaling 6 is a novel suppressor of breast tumor initiation and progression. Carcinogenesis 34, 1747–1755 (2013).

22. Yang, J., et al. RGS6 is an essential tumor suppressor that prevents bladder carcinogenesis by promoting p53 activation and DNMT1 downregulation. Oncotarget 7, 69159–69172 (2016).

23. Chatterjee, T.K., Liu, Z. & Fisher, R.A. Human RGS6 gene structure, complex alternative splicing, and role of N terminus and G protein gamma-subunit-like (GGL) domain in subcellular localization of RGS6 splice variants. J Biol Chem 278, 30261–30271 (2003).

24. Chograni, M., Alkuraya, F.S., Maazoul, F., Lariani, I. & Chaabouni-Bouhamed, H. RGS6: a novel gene associated with congenital cataract, mental retardation, and microcephaly in a Tunisian family. Invest Ophthalmol Vis Sci 56, 1261–1266 (2014).

25. Ahlers-Dannen, K.E., et al. Protein Profiling of RGS6, a Pleiotropic Gene Implicated in Numerous Neuropsychiatric Disorders, Reveals Multi-Isoformic Expression and a Novel Brain-Specific Isoform. eNeuro 9(2022).

26. Yang, J., et al. RGS6, a modulator of parasympathetic activation in heart. Circ Res 107, 1345–1349 (2010).

27. Zhang, S., Coso, O.A., Lee, C., Gutkind, J.S. & Simonds, W.F. Selective activation of effector pathways by brain-specific G protein beta5. J Biol Chem 271, 33575–33579 (1996).

28. Saunders, A., et al. Molecular Diversity and Specializations among the Cells of the Adult Mouse Brain. Cell 174, 1015–1030 e1016 (2018).

29. Chen, M., et al. GPS 6.0: an updated server for prediction of kinase-specific phosphorylation sites in proteins. Nucleic Acids Res 51, W243–W250 (2023).

30. Schildge, S., Bohrer, C., Beck, K. & Schachtrup, C. Isolation and culture of mouse cortical astrocytes. J Vis Exp (2013).

31. Ahlers-Dannen, K.E., et al. A splice acceptor variant in RGS6 associated with intellectual disability, microcephaly, and cataracts disproportionately promotes expression of a subset of RGS6 isoforms. J Hum Genet 69, 145–152 (2024).

32. Snow, B.E., Betts, L., Mangion, J., Sondek, J. & Siderovski, D.P. Fidelity of G protein beta-subunit association by the G protein gamma-subunit-like domains of RGS6, RGS7, and RGS11. Proc Natl Acad Sci U S A 96, 6489–6494 (1999).

33. Ezkurdia, I., et al. Most highly expressed protein-coding genes have a single dominant isoform. J Proteome Res 14, 1880–1887 (2015).

34. Huang, J., et al. RGS6 suppresses Ras-induced cellular transformation by facilitating Tip60-mediated Dnmt1 degradation and promoting apoptosis. Oncogene (2013).

35. Chatterjee, T.K., Eapen, A.K. & Fisher, R.A. A truncated form of RGS3 negatively regulates G protein-coupled receptor stimulation of adenylyl cyclase and phosphoinositide phospholipase C. J Biol Chem 272, 15481–15487 (1997).

36. Dulin, N.O., et al. Regulator of G protein signaling RGS3T is localized to the nucleus and induces apoptosis. J Biol Chem 275, 21317–21323 (2000).

37. Mittmann, C., et al. Evidence for a short form of RGS3 preferentially expressed in the human heart. Naunyn Schmiedebergs Arch Pharmacol 363, 456–463 (2001).

38. Zhang, K., et al. Structure, alternative splicing, and expression of the human RGS9 gene. Gene 240, 23–34 (1999).

39. Snow, B.E., et al. GTPase activating specificity of RGS12 and binding specificity of an alternatively spliced PDZ (PSD-95/Dlg/ZO-1) domain. J Biol Chem 273, 17749–17755 (1998).

40. Chatterjee, T.K. & Fisher, R.A. Novel alternative splicing and nuclear localization of human RGS12 gene products. J Biol Chem 275, 29660–29671 (2000).

41. Anderson, G.R., et al. Expression and localization of RGS9-2/G 5/R7BP complex in vivo is set by dynamic control of its constitutive degradation by cellular cysteine proteases. J Neurosci 27, 14117–14127 (2007).

42. Anderson, G.R., et al. R7BP complexes with RGS9-2 and RGS7 in the striatum differentially control motor learning and locomotor responses to cocaine. Neuropsychopharmacology 35, 1040–1050 (2010).

43. Posokhova, E., Uversky, V. & Martemyanov, K.A. Proteomic identification of Hsc70 as a mediator of RGS9-2 degradation by in vivo interactome analysis. J Proteome Res 9, 1510–1521 (2010).

44. Ostrovskaya, O.I., et al. Inhibitory Signaling to Ion Channels in Hippocampal Neurons Is Differentially Regulated by Alternative Macromolecular Complexes of RGS7. J Neurosci 38, 10002–10015 (2018).

45. Muntean, B.S. & Martemyanov, K.A. Association with the Plasma Membrane Is Sufficient for Potentiating Catalytic Activity of Regulators of G Protein Signaling (RGS) Proteins of the R7 Subfamily. J Biol Chem 291, 7195–7204 (2016).

46. Ostrovskaya, O., et al. RGS7/Gbeta5/R7BP complex regulates synaptic plasticity and memory by modulating hippocampal GABABR-GIRK signaling. Elife 3, e02053 (2014).

47. Zhou, H., et al. GIRK channel modulation by assembly with allosterically regulated RGS proteins. Proc Natl Acad Sci U S A 109, 19977–19982 (2012).

48. Kaiser, J., et al. Convergence on CaMK4: A Key Modulator of Autism-Associated Signaling Pathways in Neurons. Biol Psychiatry 97, 439–449 (2025).

49. Liu, Z., Chatterjee, T.K. & Fisher, R.A. RGS6 interacts with SCG10 and promotes neuronal differentiation. Role of the G gamma subunit-like (GGL) domain of RGS6. J Biol Chem 277, 37832–37839 (2002).

50. Ho, D.H., Han, S.J. & Son, I. The Multifaceted Role of LRRK2 in Parkinson’s Disease. Brain Sci 15(2025).

51. Bifsha, P., Yang, J., Fisher, R.A. & Drouin, J. Rgs6 is required for adult maintenance of dopaminergic neurons in the ventral substantia nigra. PLoS Genet 10, e1004863 (2014).

52. Ahlers, K.E., Chakravarti, B. & Fisher, R.A. RGS6 as a Novel Therapeutic Target in CNS Diseases and Cancer. AAPS J 18, 560–572 (2016).

53. Stewart, A., Maity, B. & Fisher, R.A. Two for the Price of One: G Protein-Dependent and - Independent Functions of RGS6 In Vivo. Prog Mol Biol Transl Sci 133, 123–151 (2015).

