## Supplemental data for "Molecular cloning of a novel, nervous system-specific RGS6 isoform lacking canonical G protein regulatory effects and with dominant negative actions"

**Data Supplement**

**Supplemental Table 1** – RGS6 specific primers.

| **Primer Name** | **Forward or Reverse Primer** | **Primer Sequence (5’-3’)** |
| --- | --- | --- |
| RGS6L-5’-H | Forward | GATACTTCCAGTCTTCCGATGTTGTGATC |
| RGS6L-ATG-H | Forward | ATGGCTCAAGGATCCGGGGA |
| RGS6L-ATG-M | Forward | ATGGCTCAGGGGTCCGGGGA |
| RGS6L-ATG-H-KpnI | Forward | AAAGGTACCTCAGGATGGCTCAAGGATCCGG |
| RGS6-18-H | Forward | CCAGAAAGTGAGCAAGGTCGTA |
| RGS6-A3R-H | Reverse | CCTCGGGGGTGAAATGCAGCG |
| RGS6-A3R-M | Reverse | CCTCGGGGGTGAAACGCGGCG |
| RGS6-αR-H-NotI | Reverse | TTTGCGGCCGCGTCAGGAGGACTGCATCAGG |
| RGS6-αR2-H | Reverse | TTCATTGCCCTCACTCTTTCTCCT |
| RGS6-αR2-H-NotI | Reverse | TTTGCGGCCGCTTCATTGCCCTCACTCTTTCTCCT |
| RGS6-βR2-H | Reverse | ACGACGTGTTCTCCCCTGAAT |
| RGS6-βR3-H | Reverse | TGCTGAAGATGTGATTTGGACATTTTA |
| RGS6-βR4-H | Reverse | AATGAGCTGGGATCAGGGCC |
| RGS6-3’-θ-H | Reverse | TGCTCCATCTCCATTTCCCAGTG |
| RGS6-A1R-H | Reverse | TCAGGCCATCATGGAGTGAAGG |
| RGS6-A2R-H | Reverse | CCTTGACCGTGACATTGGCAGT |

**Supplemental Table 2** – RGS6B shRNA and miRNA sequences

| **Construct Name** | **Sequence (5’-3’)** |
| --- | --- |
| shRNA 6B.1 | AGTCAGAATGGCCGCCGCGTTCAAGAGACGCGGCGGCCATTCTGACT |
| shRNA 6B.2 | GCAGAGTCAGAATGGCCGCTTCAAGAGAGCGGCCATTCTGACTCTGC |
| miRNA 6B | GCTGGCGCAGAGTCAGAATGCTGTAAAGCCACAGATGGGTATTCTGACTCTGCGCCAGC |

**Supplemental Table 3** – Summary of pathological findings from the University of Iowa Department of Pathology for all glioma tumor samples analyzed in Fig. 8A. Expression of both RGS6B and RGS6L are categorized as Low = <20%, Reduced = <75%, Elevated = > 120%, or unchanged (-) relative to non-malignant control samples. Samples where aberrant migration of the 69 kDa RGS6B band were noted are indicated with a (*). This phenomenon was found in 50% of all samples but was more common in high grade glioblastomas (6/8).

| **Sample Number** | **Tumor Type** | **Stage** | **% Cont.**  **69 kDa** | **% Cont.**  **56 kDa** | **Aberrant Migration of**  **69 kDa Band** |
| --- | --- | --- | --- | --- | --- |
| 314 | Astrocytoma | 2 | Low | Low |  |
| 206 | Anaplastic Astrocytoma | 3 | Reduced | Reduced |  |
| 329 | Glioblastoma | 4 | Low | Low |  |
| 511 | Astrocytoma | 2 | Low | Low |  |
| 305 | Anaplastic Astrocytoma | 3 | Low | Reduced | * |
| 335 | Glioblastoma | 4 | Reduced | Reduced | * |
| 342 | Glioblastoma | 4 | Elevated | Low | * |
| 549 | Astrocytoma | 2 | - | - |  |
| 340 | Anaplastic Astrocytoma | 3 | Reduced | Reduced |  |
| 379 | Glioblastoma | 4 | Reduced | Reduced | * |
| 396 | Glioblastoma | 4 | Reduced | Reduced | * |
| 416 | Glioblastoma | 4 | Reduced | Low | * |
| 417 | Glioblastoma | 4 | Reduced | Reduced | * |
| 428 | Glioblastoma | 4 | - | - |  |
| 168 | Oligodendroglioma | 2 | Low | Low |  |
| 288 | Anaplastic Oligodendroglioma | 3 | Low | Low | * |
| 267 | Oligodendroglioma | 2 | Low | Low | * |
| 304 | Anaplastic Oligodendroglioma | 3 | - | Reduced | * |
| 647 | Oligodendroglioma | 2 | Reduced | - |  |
| 600 | Anaplastic Oligodendroglioma | 3 | - | - |  |

**Supplemental Table 4** – Exon sequences (mRNA) for newly identified RGS6 exons.

| **Exon #** | **bp** | **Exon Sequence** |
| --- | --- | --- |
| **S1** | 105 | ATGCCTGTGCCATGCTCTTGGACTTCCCAGCCTCCAGAACTGTGAGAAATAAATTTCTTTATAAATTACTGAGCATTCTATTATAGCAGCACAGACTAAGACAGT |
| **S2** | 133 | GTTTCTCAAAGGGCTGAACTGACTCAAATATACTGGGGACTCCCTCTGCTCCTCACACCTGCCCAGGAGGGTTACTTCTCAAGGTCTTGTGCCAGCCATTAGTGTTGTCCACTCAAACTGCTCAGGAAGCGTG |
| **S3** | 136 | TGGCTGGAACCTGTATCTGGTGTATCACAACGCTCCATTTCGCTGATCACACACAGCCAGAAGTGACAGTCGCTTCTCTCACTGCCAAGGACAGAGGAGACTCTGGCTGGACAGGAGAAATTCAGGTCGAACCAAG |
| **S4** | 126 | AGCAGATGTAGAGGACCAGGAGTGATTCAGAGATGAGAACTGAATCTTCCCTCCAGAAAGTTATGTTCCAGAGGGAAAGAAGCGGAACTTCAGTAACAGAGAATATGAAATTGGTCCTAAGTAGAG |
| **A1** | 151 | AATGTGCAGAACACAAGGCCTGCGGGTGCCTGGTCTATCTTCATGGCTGCTTTGCCAATGGCCGTGTGGCTCCGCACACCAAGAAGCTGCCACGGGCTGTCAGAGTCAAGGCTTCTGGGACTTCCTTCACTCCATGATGGCCTGAGAGAAG |
| **A2** | 211 | GACAGAGCCTGCCTCAGTCCTGTCCGGGGAGCAGCAGCACCGCAGCAGAGGGGGCCAGAGAAAGAGCATCCAATGGCAGGAGTCTGGCAAGGCGACAGCACTGCCAATGTCACGGTCAAGGAGAGGAGACCCAACTCTGCCTGTGGTCCAGACTGGAAAGTTCAAAGAGAAGTCAGAGAGCTTCAAGTCTGAGGACTTCAAAGGCTAAGAA |
| **A3** | 67 | TGCTGCTGGCGCAGAGTCAGAATGGCCGCTGCATTTCACCCCCGAGGTGGCTTCCTTCCAGCAGTGG |
| **θ** | 47 | GAAAAGACAGACTGTCAAGCAAGGACCGAAAGCCTGGCCTTAACATC |
| **αΔ129β** | 217 | GGAAAGTCGCTGGCGGGCAAGCGCCTCACGGGCCTGATGCAGTCCTCCTGACCGTTCCTACCGCAGGTCCAGGGCCTGGGCCCGCGGACCCCACAGGCAGGCGGCGGCGCTCCACATCTGCGGACAGAGTTTCCTTACGAGGAGACTTGGTCACTGTGAAGGAGAAAGATTGTATTCCAACACTCCACTCGCTAAGAGGCCCTGATCCCAGCTCATT |

**Supplemental Table 5** – Summary of novel RGS6 mRNA sequences identified

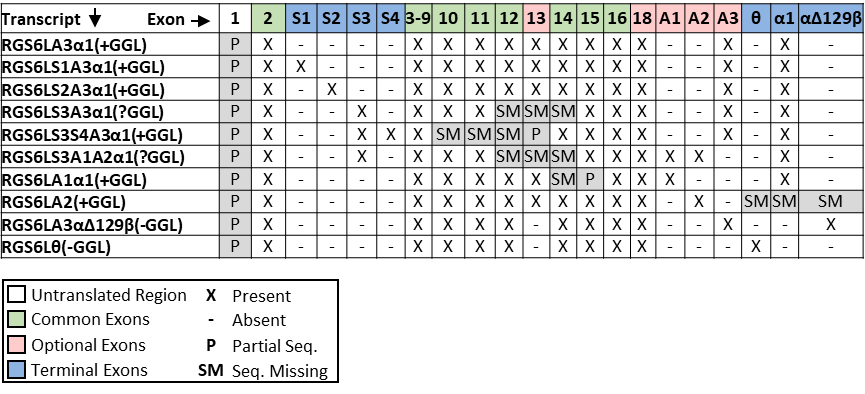

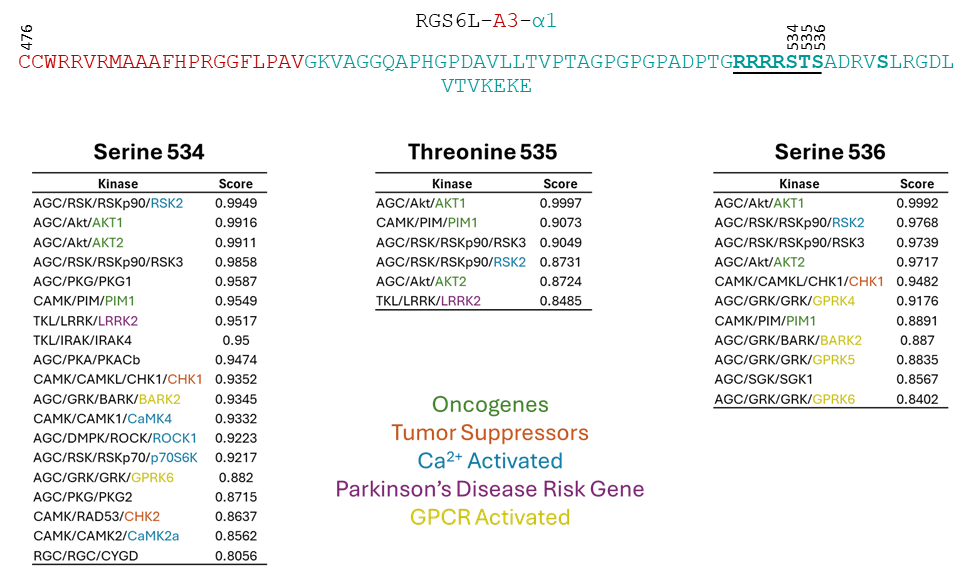

**Supplemental Figure 1** – Summary of phosphorylation site analysis for RGS6B (RGS6LA3α1). Putative phosphorylation sites identified are noted in bold text in the protein sequence of RGS6B (top). Putative RGS6B kinases identified by the GPS 6.0 webserver are noted for S534, S535, and S536, which displayed the strongest prediction scores (>0.80). Functional relevant kinase candidates are color-coded for easy identification.

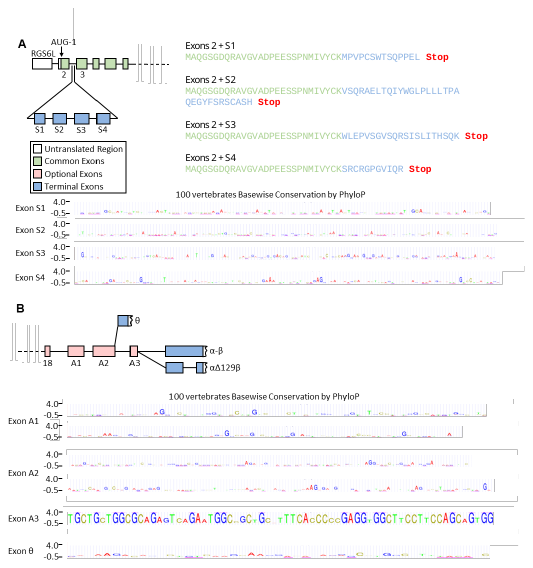

**Supplemental Figure 2** – Conservation of novel exon sequences across species. Summary of species conservation was generated from 100 vertebrates by PhyloP for (A) the novel S1-S4 stop exons and (B) novel exons A1, A2, A3, and θ.

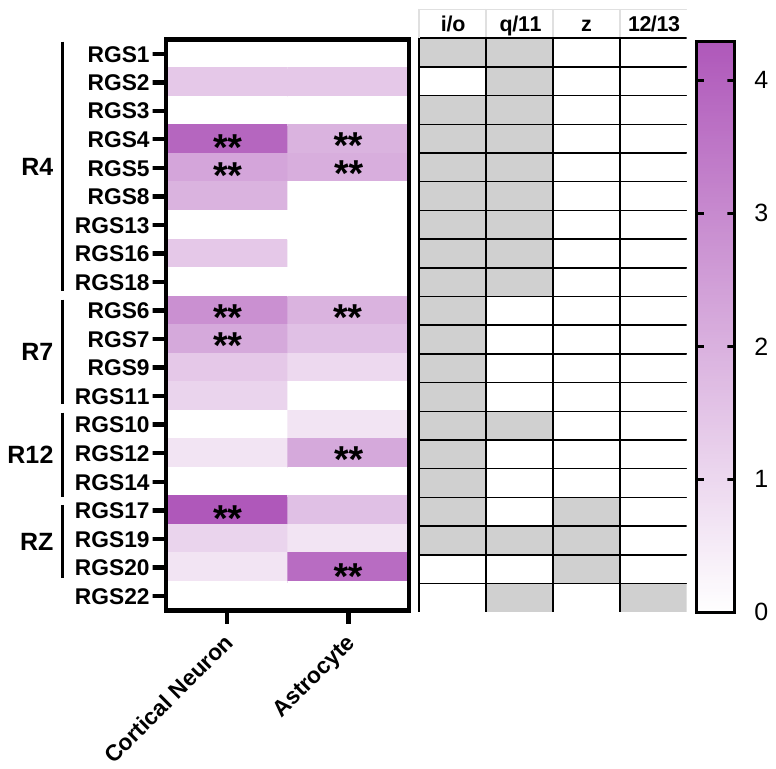
**Supplemental Figure 3** – RGS protein expression in cell types of the frontal cortex was extracted from a single cell nuclear RNA seq (scRNAseq) cell atlas of the murine mouse brain available online at Dropviz.org.** indicate the 5 RGS proteins with highest expression in each cell type.

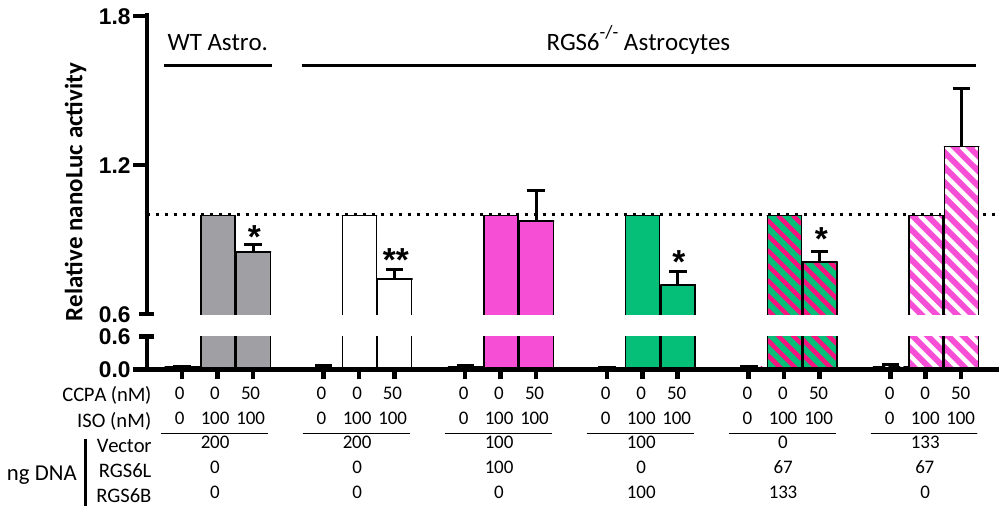

**Supplemental Figure 4** – RGS6^-/-^ or WT astrocytes were co-transfected with a dual luciferase reporter DNA in addition of either RGS6L, RGS6B, or control DNA for 48 hrs. Cells were then treated with control vehicle (DMSO) or a cocktail of isoproterenol with various concentrations of CCPA (A1 receptor agonist). Luciferase was used to evaluate the ability of RGS6 isoforms to inhibit CCPA-induced activation of Gα_i_, via measuring p-CREB-mediated activation of CRE promoter in the luciferase reporter construct. Results are means ± S.E.M. of three experiments. *, P<0.05; **, P<0.01.
